# Selectivity and engineering of the sialoglycan-binding spectrum in Siglec-like adhesins

**DOI:** 10.1101/796912

**Authors:** Barbara A. Bensing, Lioudmila V. Loukachevitch, Rupesh Agarwal, Izumi Yamakawa, Kelvin Luong, Azadeh Hadadianpour, Hai Yu, Kevin P. Fialkowski, Manuel A Castro, Zdzislaw Wawrzak, Xi Chen, Jerome Baudry, Jeremy C. Smith, Paul M Sullam, T M Iverson

## Abstract

The Siglec-like Serine-Rich Repeat (SRR) adhesins mediate bacterial attachment to mammalian hosts via sialoglycan receptors. Here, we combine structural, computational, biochemical, and phylogenetic approaches to elucidate the determinants of the sialoglycan-binding spectrum across the family of Siglec-like SRR adhesins. We further identified mutable positions that disproportionately affect sialoglycan selectivity, as demonstrated by increases in binding to alternative ligands of 2- to 3-orders of magnitude. Biologically, these studies highlight how bacteria nimbly modulate the receptor interaction during coevolution of host and pathogen. These studies additionally created binding proteins specific for sialyl-T antigen or 6S-sialyl Lewis^X^ that can recognize glycosylation of human plasma proteins. The engineered binding proteins can facilitate the characterization of normal cellular glycan modifications or may be used as diagnostic tools in disease states with altered glycosylation.

**Significance:** The ability of bacteria to bind selectively to host receptors underlies both commensalism and pathogenesis. Here, we identify the molecular basis for receptor selectivity in streptococci that bind to sialoglycan receptors. This revealed how to convert these adhesins into selective probes that measure triand tetrasacharides within the context of larger glycosylations. These probes that can be used in a laboratory with no specialized equipment and can be used to address biological questions relating to sialoglycan-dependent signaling and adhesion.

---

The decoration of proteins with sialoglycans is functionally important in numerous mammalian signaling pathways. However, a wide array of bacterial and viral adhesive proteins exploit these sialoglycans as host receptors during infection. In these cases, sialoglycan selectivity determines whether a pathogen can adhere to a preferred anatomical niche or can infect a particular host (1, 2).

<u>S</u>ialic acid binding immuno<u>g</u>lobulin like <u>lec</u>tin (Siglec)-like adhesins are found within the larger family of Serine-rich repeat (SRR) adhesins (3-14), which form fibril-like protrusions on streptococci and staphylococci (15). The Siglec-like adhesins (3, 7, 16-23) always include two adjacent modules: a “Siglec” domain and a “Unique” domain (21). In oral streptococci, Siglec-like adhesins bind to carbohydrates containing a terminal Siaα2-3Gal (Sia = Neu5Ac or Neu5Gc) (3, 4, 7-9, 16-19, 22) at a YTRY sequence motif (16, 18, 21, 24). In humans, Neu5Acα2-3Gal is commonly found at the termini of the complex *O*-linked sialoglycans that modify the MUC7 salivary mucin (9, 16, 25) or glycoproteins in both blood plasma (17) and on platelets (8, 23). Binding to α2-3-linked sialoglycans may therefore allow colonization of the oral cavity, or can lead to endovascular infection (4, 21, 22, 26).

Sequences of the Siglec-like adhesins are quite variable, as are the host sialoglycans to which they bind. For example, GspB from *Streptococcus gordonii* strain M99 binds with narrow selectivity to the sialyl-T antigen (sTa) trisaccharide (3, 16, 19) (for carbohydrate structures, see **Fig. S1**). In contrast, other SRR adhesins, such as Hsa from *S. gordonii* strain Challis, bind with high avidity to multiple glycans (3, 16, 19).

Here, we evaluated sialoglycan binding and selectivity of Siglec-like adhesins using structural, computational, and biochemical approaches. We then used this information to engineer adhesins with altered binding properties and showed that this affected the preferred host receptor. Our findings provide insights into the molecular basis for sialoglycan selectivity by Siglec-like adhesins and suggest a route for developing these adhesins into a broad array of tools to characterize sialoglycan distribution.

## Results

### Selection of representative adhesins

We began by correlating phylogenetic analysis of sialoglycan-binding Siglec and Unique domains (**Fig. S2**) with reported sialoglycan selectivity (3, 16, 17, 19, 20). This identified that evolutionary relatedness is a moderate, but not strong, predictor of glycan selectivity. In short, most of the adhesins of the first major branch of the tree (blue in **Fig. S2**) bound two or more related tri- or tetrasaccharides, albeit without a clear glycan preference (3, 16, 17, 19, 20). In contrast, the four characterized adhesins of the second major branch (green in **Fig. S2**) exhibit narrow selectivity for sTa (3, 16, 17, 19, 20).

From the first branch of the tree (blue in **Fig. S2**), we selected the Siglec and Unique domains of Hsa (termed Hsa_Siglec+Unique_), and the equivalent domains from *Streptococcus sanguinis* strain SK678 and *Streptococcus mitis* strain NCTC10712 for further study. These three adhesins are >80% identical but exhibit different receptor selectivity. Hsa_Siglec+Unique_ binds detectably to a broad range of Siaα2-3Galβ1-3/4HexNAc glycans but not to fucosylated derivatives (16, 19). In comparison, SK678_Siglec+Unique_ exhibits narrow selectivity for 3’-sialyl-*N*-acetyllactosamine (3’sLn) and 6-*O*-sulfo-sialyl Lewis X (6S-sLe^X^), while NCTC10712_Siglec+Unique_ binds strongly to a range of 3’sLn-related structures (16). The combination of high sequence identity and distinct binding spectrum suggests that we will be able to pinpoint the origins of sialoglycan selectivity with these comparators.

The second major branch of the evolutionary tree (green in **Fig. S2**) includes GspB from *S. gordonii* strain M99 (3, 8, 9, 21, 27). GspB_Siglec+Unique_ exhibits narrow selectivity for the sTa trisaccharide, as do the other previously-characterized members of this evolutionary branch (3, 16, 17, 19, 20). In seeking comparators of GspB, we performed binding studies on additional homologs. We identified that the Siglec and Unique domains of the adhesin from *S. gordonii* strain SK150 (termed SK150_Siglec+Unique_) are 62% identical to the corresponding regions of GspB but exhibit broader carbohydrate selectivity (**Fig. S3**). The distinct binding properties make these good comparators for understanding sialoglycan selectivity.

### Structures of the Hsa-like and GspB-like adhesins

Using these five comparators, we evaluated how sequence differences affect the structure. As determined by crystallography (**Fig. 1A-1D Table S1, S2**), all five adhesins exhibited similar folds of the individual domains (**Fig. 1A, 1B**). However, the interdomain angle differed between the Hsa-like and GspB-like adhesins in a way that correlates with phylogeny (**Fig. 1C, 1D, S4A**).

**Figure 1.**
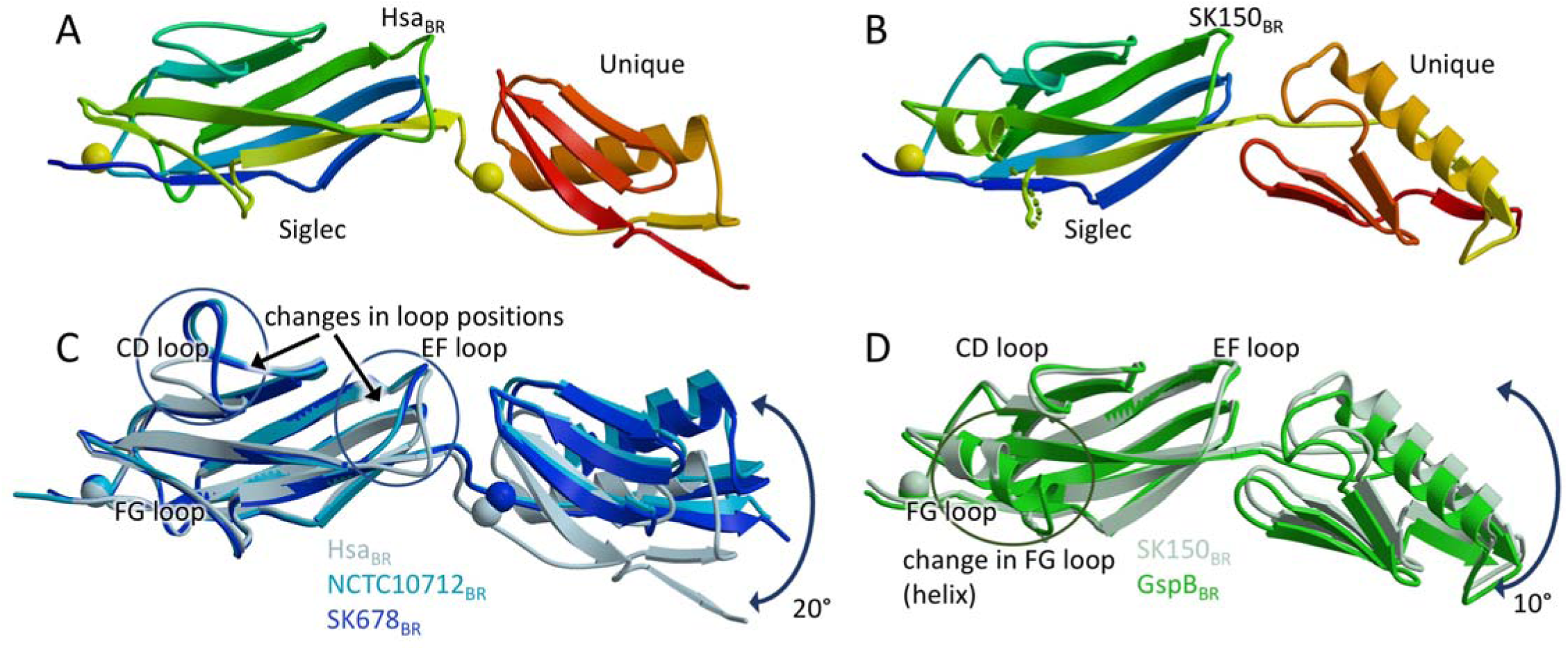
Structures of sialoglycan binding regions of Siglec-like SRR adhesins. A and B. Ribbon diagrams of A. Hsa-_Siglec_Unique_ and B. SK150_Siglec+Unique_ with the N-terminus in *blue* and the C-terminus in *red*. Ions are shown as spheres. C. Hsa-like adhesins. Hsa_Siglec+Unique_ is in *grey*, NCTC10712_Siglec+Unique_ is in *cyan*, and SK678_Siglec+Unique_ is in *blue*. D. GspB-like adhesins. GspB_Siglec+Unique_ is in *green* and SK150_Siglec+Unique_ is in *light green*.

Notably, even in closely-related adhesins, the Siglec domain contains conformational differences in three loops of the V-set Ig fold: the CD loop, the EF loop, and the FG loop (**Fig. 1C, 1D**). Sequence variation of these adhesins disproportionately maps to these loops (**Fig. S4D**). Taken together, these studies of unliganded adhesins identify features that correlate with phylogeny and reveal regions of disproportionate variation in closely related adhesins.

### Sialoglycan binding and conformational selection

We next determined costructures of sTa with Hsa_Siglec+Unique_ (**Fig. 2A**) or the Siglec domain of GspB (GspB_Siglec_) (**Fig. 2B**). In both costructures, sTa binds in a defined pocket of the Siglec domain (**Fig. 2A, 2B**). This pocket is analogous to the sTa-binding site identified in the SRR adhesin SrpA from *S. sanguinis* strain SK36 (18, 20) (**Fig. 2C**), which phylogenetically groups with Hsa (**Fig. S2**). Interactions between sTa and each adhesin involves the YTRY sialic acid-binding motif (Hsa^338-341^ or GspB^482-485^) (**Fig. 2D, Fig. S5**)(16) and three inserts of the V-set Ig fold: the CD loop (Hsa^284-296^ or GspB^440-453^), the EF loop (Hsa^330-^ 336 or GspB^475-481^), and the FG loop (Hsa^352-364^ or GspB^499-^ 511) (**Fig. 2, Fig. S5**). These same regions vary disproportionately in both sequence and conformation in the unliganded structures (**Fig. 1, S4D**).

**Figure 2.**
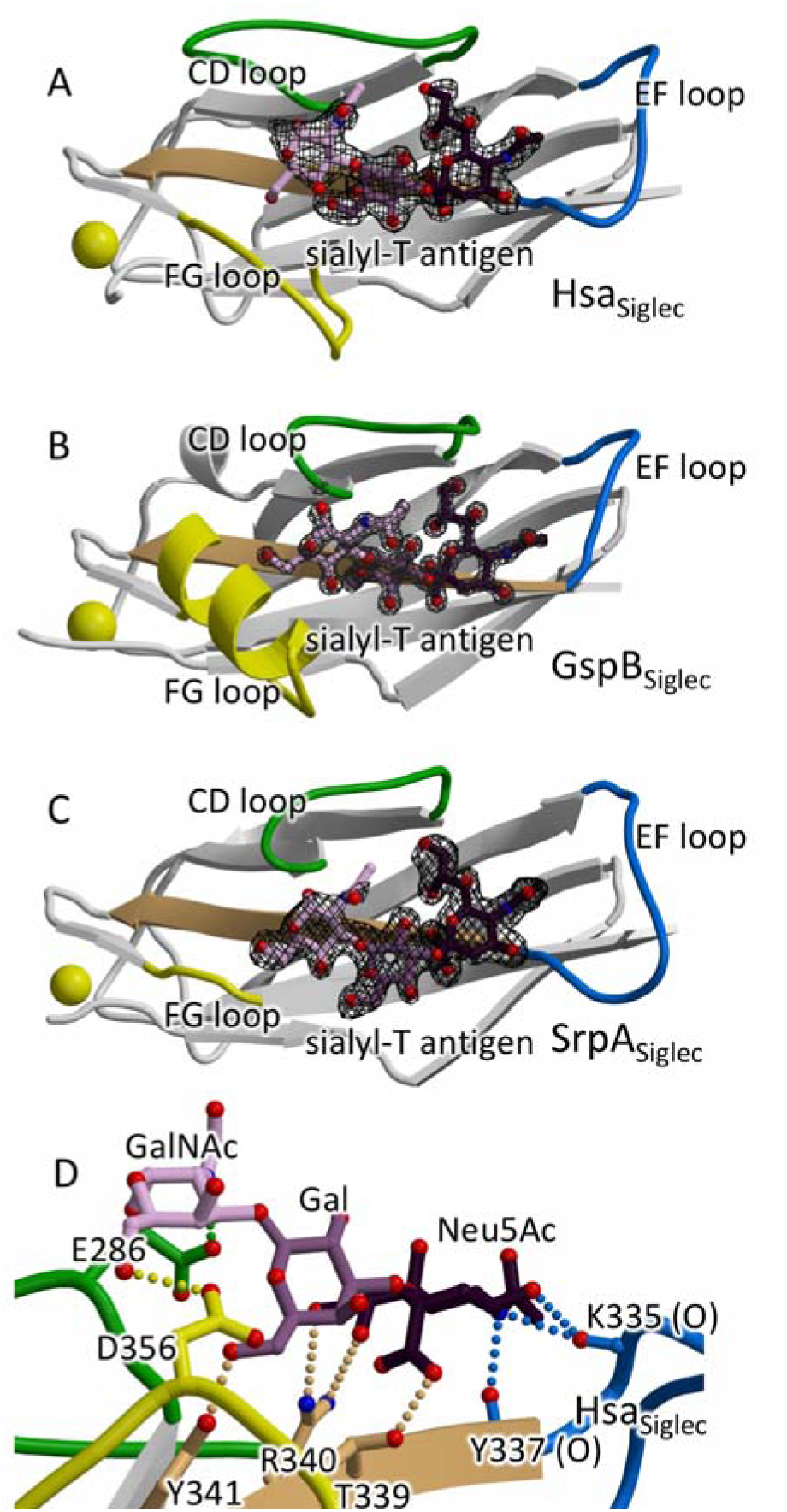
Costructures of sTa bound to Siglec-like SRR adhesins. A-C. Siglec domain of A. Hsa, B. GspB, and C. SrpA (PDB entry 5IJ3 (18)) bound to sTa. *F*_*o*_ – *F*_*c*_ omit electron density contoured at 3σ is shown as black mesh. In each panel, the F-strand harboring the “YTRY” motif is shown in *tan*, variable loops that interact with the ligand are shown in *green* (CD loop), *blue* (EF loop), and *yellow* (FG loop). D. Contacts between Hsa_Siglec_ and sTa. The color of the hydrogen-bond reflects the structural element involved in the interaction.

The YTRY motif is located on the F-strand of the V-set Ig fold and contributes to binding the invariant terminal Siaα2-3Gal of the target *O-*linked sialoglycans (18, 21, 24). However, the role of the three loops in glycan affinity and selectivity is unknown. We queried whether these loops exhibited inherent flexibility, a property believed to correlate with the ability to evolve binding to new ligands (28-30). Temperature factor analysis suggests that these loops have high flexibility in the absence of ligand (**Fig. S6**). Moreover, these loops exhibit conformational differences between the ligand-bound and ligand-free structures (**Fig. 1, 3**). In the GspB_Siglec_ structure, the helix of the FG loop rotates 10° in response to sTa which results in a maximal displacement of 1.3 Å (**Fig. 3A**) while in the Hsa_Siglec+Unique_ structure, the EF loop moves 5.9 Å (**Fig. 3B**) and allows the Hsa^K335^ carbonyl to form hydrogen-bonding interactions to the Neu5Ac C5 nitrogen and C4 hydroxyl.

**Figure 3.**
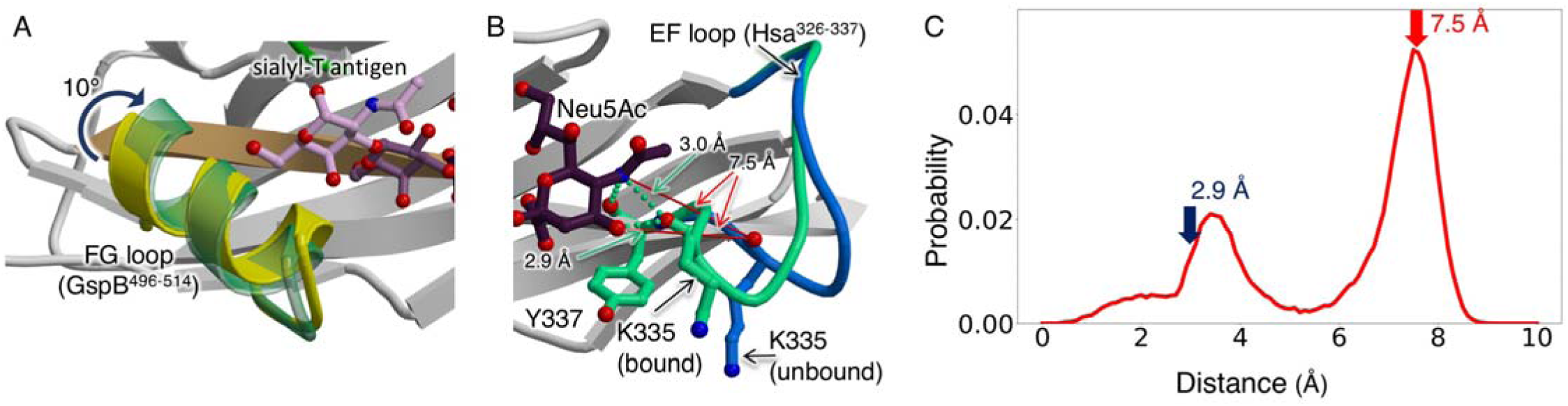
Conformational changes associated with sTa binding. A. The FG loop of GspB_Siglec_ rotates 10° upon sTa binding. B. The EF loop of Hsa_Siglec+Unique_ adjusts to promote formation of hydrogen-bonding interactions between Hsa^K335^ and the Neu5Ac of sTa. C. Probability of distance distribution between the Neu5Ac O4 hydroxyl of sTa and the Hsa^K335^ backbone carbonyl, as calculated by MD simulations. A bimodal distribution of distances exhibit maxima at 7.5 Å, which reflects the unliganded crystal structure, and at 3.5 Å, which approaches the liganded crystal structure. The formation of the hydrogen-bond between the Hsa^K335^ carbonyl and Neu5Ac likely further shifts the conformational equilibrium to a pose that supports the 2.9 Å distance observed in the bound state.

To explore the conformations available to these loops, we performed MD simulations of unliganded Hsa_Siglec+Unique_ and GspB_Siglec+Unique_. The loops surrounding the glycan binding pocket exhibited considerably more flexibility than other parts of the protein (**Fig. S7A–D**). Moreover, the ligand-bound conformation is among the predicted conformations sampled in the absence of ligand (**Fig. 3, S7**). Of particular note is the main chain carbonyl of Hsa^K335^, which forms a hydrogen bond to sTa in the experimental costructure and samples both the bound and unbound states in the apo form (**Fig. 3B, 3C**). These calculations predict that sTa shifts the equilibrium of the EF loop to the position observed in the crystal structure of the bound state (**Fig. 2D, 3B, 3C, S5A**). Together, these analyses support a conformational selection mechanism over an induced fit mechanism, a property that may allow adaptation to changes of the host *O*-glycan receptors.

To experimentally assess whether conformational selection could contribute to ligand binding, we focused on the broadly selective Hsa_Siglec+Unique_. We introduced rigidifying prolines or replaced glycines at predicted hinges (Hsa^N333P^, Hsa^G287A/G288P^), both of which are predicted to reduce the flexibility required for conformational selection. As controls, we developed variants that introduced glycines (Hsa^L363G^, Hsa^S253G^) (**Fig. S8A**). Hsa^N333P^ was associated with substantially reduced sialoglycan binding for all ligands tested; Hsa^G287A/G288P^ also exhibited reduced binding, but the effect was less pronounced (**Fig. S8B – S8D**). In contrast, glycine-substituted Hsa^L363G^ and Hsa^S253G^ exhibited binding similar to wild-type (**Fig. S8B – S8D**). These experiments provide support for a conformational selection mechanism.

### Sialoglycan binding spectrum

All characterized ligands of the Siglec-like SRR adhesins contain a Siaα2-3Gal disaccharide at the non-reducing terminus (16, 19). However, the identity of, and linkage to, the adjacent sub-terminal sugar varies. Analysis of the contacts in the costructures of Hsa_Siglec+Unique_ and GspB_Siglec_ with sTa identified that the sub-terminal sugar predominantly contacts the CD loop and the FG loop of the Siglec domain (**Fig. 2, S5**). In contrast, the Neu5Acα2-3Gal interacts with the YTRY motif and residues in the EF loop (**Fig. 2, S5**).

To test how these loops affect sialoglycan selectivity, we engineered chimeras with the backbone of one adhesin and the loops of a closely-related adhesin. In the SK678^Hsa-loops^ and NCTC10712^Hsa-loops^ chimeras, selectivity became more similar to that of Hsa than the parent adhesin (**Fig. 4, Table S3**). This indicates that a major determinant of selectivity in Hsa-like adhesins is the combined contribution of the CD, EF, and FG loops. We next assessed whether one loop dominates this effect using individual substitutions. SK678^Hsa-CD-loop^ exhibited substantially decreased affinity for 3’sLn and 6S-sLe^X^, while SK678^Hsa-EF-loop^ had somewhat increased binding for all of the glycans tested, and SK678^Hsa-FG-loop^ had moderately decreased binding for 3’sLn and a substantially decreased affinity for 6S-sLe^X^ (**Fig. S9A**). NCTC10712^Hsa-CD-loop^ and NCTC10712^Hsa-FG-loop^ exhibited differential changes in binding for the sialoglycan ligands, while NCTC10712^Hsa-EF-loop^ increased the range of bound ligands, but left binding unchanged for the preferred ligands (**Fig. S9B**).

**Figure 4.**
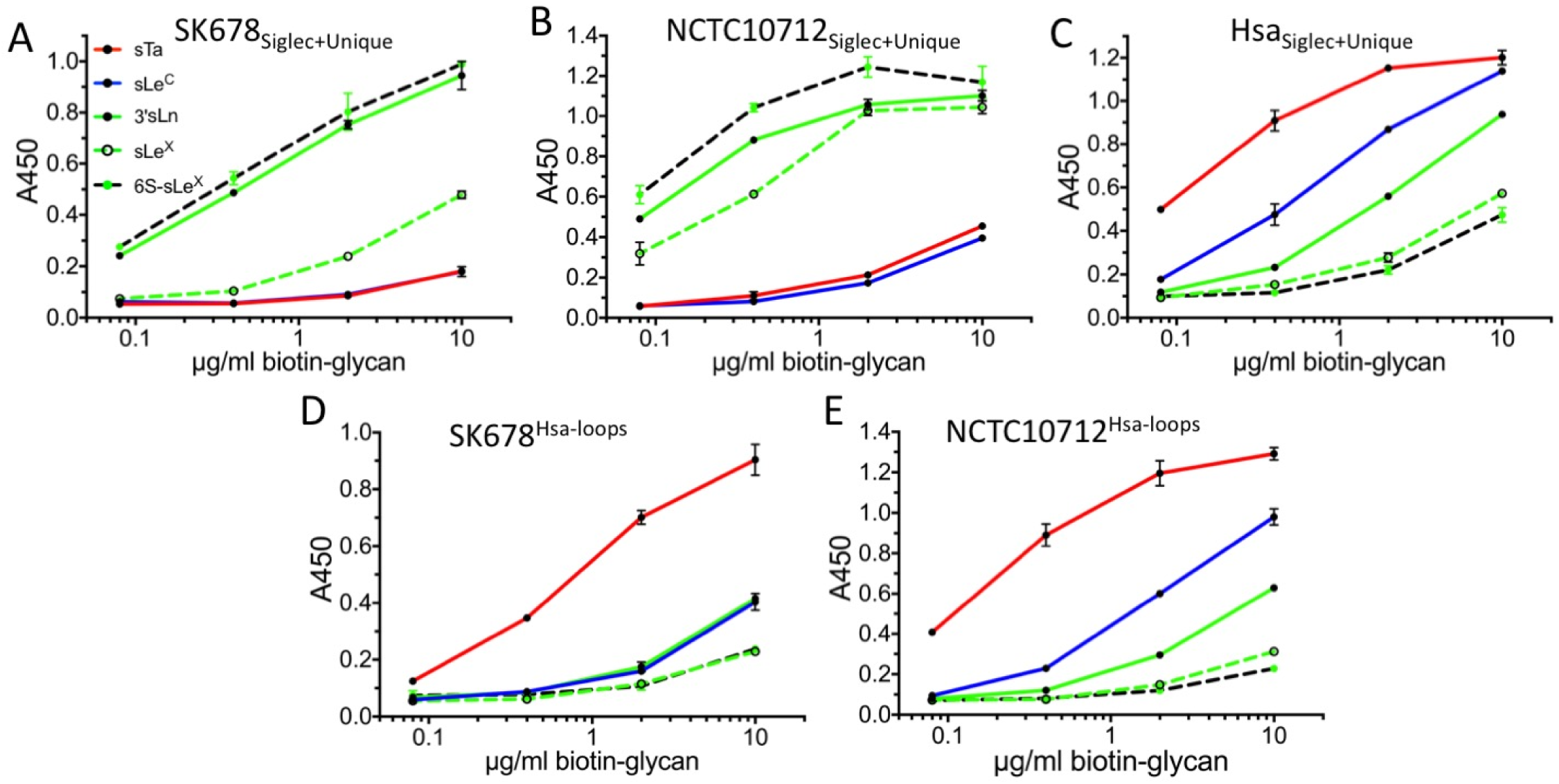
Chimeragenesis of Hsa-like adhesins. Dose-response curves of A. wild-type GST-SK678_Siglec+Unique_, B. wild-type GST-NCTC10712_Siglec+Unique_, and C. wild-type GST-Hsa_Siglec+Unique_ to five selected ligands. D and E. Dose-response curves of the chimeras D. GST-SK678^Hsa-loops^ and E. NCTC10712^Hsa-loops^ which contain the CD, EF, and FG loops of Hsa. In each case, sTa binding increases. Measurements were performed using 500 nM of immobilized GST-adhesin and the indicated concentrations of each ligand, and are shown as the mean±SD (n = 2).

One interpretation of the chimeragenesis data takes into consideration the position of each loop with respect to the ligand (**Fig. 2, S5**). The residues of the EF loop only interact with sialic acid (**Fig. S5**) and may act in concert with the YTRY motif to support binding of the invariant region of the ligands, i.e. Siaα2-3Gal. Yet substitution of the EF loop of the more promiscuous Hsa into SK678 and NCTC10712 resulted in a somewhat broader binding spectrum (**Fig. S9A, S9B**). We posit that flexibility of the EF loop (**Fig. 3B, 3C, S6, S7**) adjusts the orientation of the entire sialoglycan to optimize the interaction between the variant position of the ligand and the CD and FG loops (**Fig. 3**). If the EF loop controls the ligand orientation, then the CD and FG loops may act in synergy to select the glycan. In particular, the FG loop of Hsa restricts the binding pocket and inhibits accommodation of Fucα1-3GlcNAc, as reflected by the lower binding of sLe^X^ and 6S-sLe^X^ to SK678^Hsa-FG-loop^ and NCTC10712^Hsa-FG-loop^ (**Fig. 5D, 5E, S7A, S7B)**.

**Figure 5.**
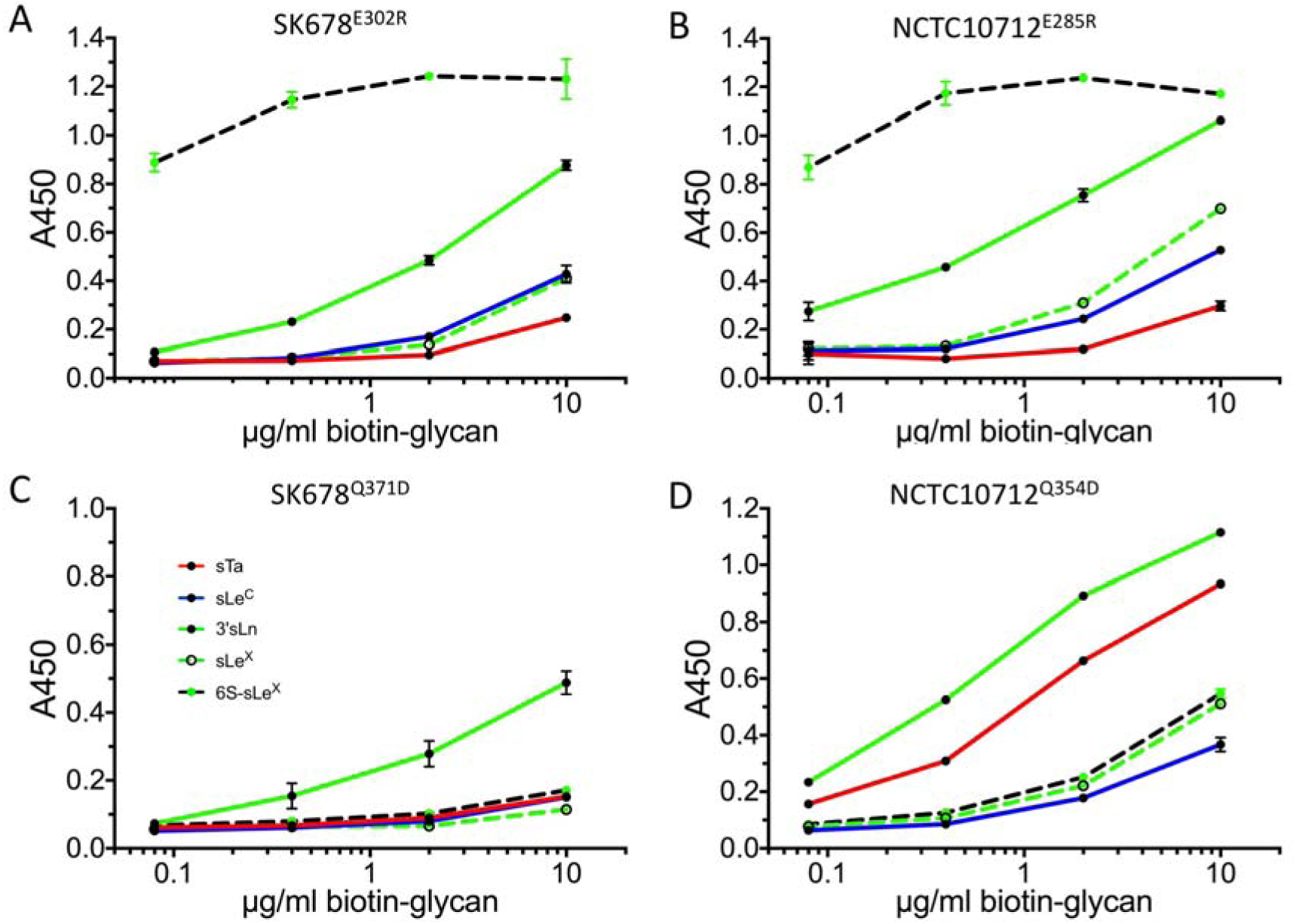
Binding selectivity of select engineered adhesins. Dose response curves of A. GST-SK678^E302R^, B. GST-NCTC10712^E285R^, C. GST-SK678^Q371D^, and D. GST-NCTC10712^Q345D^. When compared to wild-type (see **Fig. 5A, 5B**), the GST-SK678^E302R^ and GST-NCTC10712^E285R^ variants exhibit increased binding to 6S-sLe^X^, and reduced binding to 3’sLn and sLe^X^. Conversely, both the GST-SK678^Q371D^ and GST-NCTC10712^Q345D^ variants have substantially reduced binding to the fucosylated ligands sLe^X^ and 6S-sLe^X^. Measurements were performed using 500 nM of immobilized GST-adhesin and the indicated concentrations of each ligand, and are shown as the mean±SD (n = 2).

We next evaluated chimeras of the GspB-like adhesins. GspB^SK150-CD-loop^ and GspB^SK150-FG-loop^ substantially decreased glycan affinity; as with the Hsa-like adhesins, GspB^SK150-EF-loop^ had little impact (**Fig. S9C**). However, in the GspB^SK150-loops^ chimera, which substituted all three loops, the binding affinity remained low (**Fig. S9C**). One explanation for the uneven success of chimeragenesis is that the Hsa-like chimeras used starting adhesins with more flexible loops that could better adjust to the non-native scaffold. It is also possible that the Hsa-like adhesins benefitted from a better starting match between the sequences. To evaluate these possibilities, we engineered GspB-SK150 “mini-chimeras” swapping only residues that directly contact the ligand (**Fig. S5**). The GspB^L442Y/Y443N^ mini-chimera had increased binding to 3’sLn and sLe^c^ and was overall more similar in selectivity to SK150 than to GspB (**Fig. S3, S10A, S10B**); the converse mini-chimera of SK150 still exhibited reduced binding (**Fig. S10C**). The incomplete success of the mini-chimeras suggests that both the sequence match of the starting adhesins and the limited loop flexibility impacted the ability to alter selectivity via chimeragenesis.

### Engineered adhesins with altered selectivity

If we are correct that selectivity is largely conferred by the CD and FG loops, we should be able to engineer the binding spectrum through mutation of these loops. We selected the Hsa-like adhesins, where chimeragenesis had greater success (**Fig. 4**), possibly as a result of increased loop flexibility. Hsa^E286^ (in the CD loop) and Hsa^D356^ (in the FG loop) each directly contact the GalNAc of sTa (**Fig. 2D)**. In these positions and the equivalent positions of SK678 and NCTC10712, we substituted residues predicted to alter hydrogen-bonding characteristics.

As measured by ELISA, our engineered adhesins exhibited altered binding spectra (**Table S3, Fig. 5, Fig. S11**). The most striking results were for variants of the CD loop of NCTC10712_Siglec+Unique_ and SK678_Siglec+Unique_, which become highly selective for 6S-sLe^X^ via a substantial increase in binding for this sulfated tetrasaccharide and a decrease in binding to other glycans (**Fig. 5A, 5B**). Variants of the FG loop lost binding to fucosylated ligands but had little increase in binding to alternative ligands (**Fig. 5C, 5D**). As a result, the NCTC10712^Q345D^ variant became more selective for 3’sLn while the SK678^Q371D^ variant exhibited low binding to all tested ligands. The observed loss of binding to the fucose-containing sLe^X^ and 6S-sLe^X^ by FG loop variants is consistent with the chimeragenesis showing that the FG loop is particularly important for accommodation of fucosylated sialoglycans (**Fig. S9A, S9B**).

We also found that Hsa^E286R^, Hsa^D356R^, and Hsa^D356Q^ (**Fig. S11**) increased binding to 3’sLn, sLe^C^, sLe^X^, and 6S-sLe^X^ as compared to wild-type (**Fig. 4A**), but showed different degrees of discrimination between 3’sLn, sLe^C^, and sTa. Hsa^E286R^ showed similar binding to all ligands tested and thus is even more broadly selective than wild-type (**Fig. S11A**). In contrast, Hsa^D356R^ and Hsa^D356Q^ each had an increase in 3’sLn, sLe^X^, and 6S-sLe^X^ binding, but a decrease in sTa binding. Or in other words, these variants bind to a broad range of ligands with a distinct ligand preference from wild-type via a gain-of-function mechanism.

### Engineered adhesins show differential recognition of human plasma glycoproteins

We used these engineered adhesins to assess glycosylation of plasma proteins. Our prior studies identified that Hsa_Siglec+Unique_ preferentially binds proteoglycan 4 (460 kD) in human plasma while NCTC10712_Siglec+Unique_ binds GPIbα (150 kD). These adhesins also bind different glycoforms of the C1-esterase inhibitor (100 kD) (17). Here, Far Western analysis showed that the NCTC10712^Hsa-loops^ and SK678^Hsa-loops^ chimeras recognized proteoglycan 4 rather than the preferred receptors for wild-type SK678_Siglec+Unique_ and NCTC10712_Siglec+Unique_ (**Fig. 6A**). We also found that the 6S-sLe^X^-selective SK678^E302R^ variant binds both GPIbα and the C1-esterase inhibitor (**Fig. 6B**), a binding pattern similar to that of wild-type NCTC10712_Siglec+Unique_ (**Fig. 6A**). This latter finding suggests that 6S-sLe^X^ is present on both GPIbα and C1-esterase inhibitor. It also identifies that SK678^E302R^ will be useful as a probe for detecting this modification.

**Figure 6.**
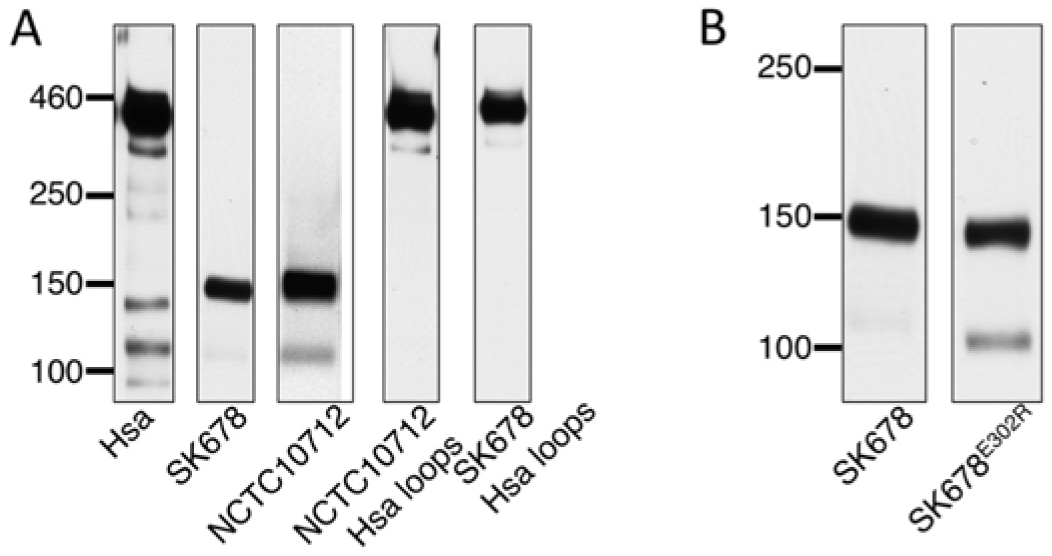
Far Western analysis of wild-type, chimeric, and engineered Hsa-like SRR adhesins binding to plasma proteins. A. Hsa-like chimeras. B. The SK678^E302R^ point mutant. As previously identified by affinity capture and mass spectrometry (17), the 460 kD band is proteoglyan 4, the 150 kD band is GP1bα, and the 100 kD band is C1-esterase inhibitor. Each Far Western was performed at least twice, with a representative blot shown.

## Discussion

Individual Siglec-like adhesins recognize sialoglycans with as few as three and possibly more than six linked sugars (16, 17, 19, 20). Many of these adhesins bind to a preferred ligand with narrow selectivity, and many, like Hsa, bind strongly to multiple ligands (16, 17, 19, 20). Our results suggest that for the Siglec-like adhesins that recognize trisaccharides, the binding pockets contain two distinct recognition regions. The first region interacts with the sialic acid-containing non-reducing terminus of the sialoglycan, i.e. Siaα2-3Gal (18, 20). This region is formed from both the YTRY motif on the F-strand (16, 18, 21, 24) and the EF loop (**Fig. 2**). The second region selects for the reducing end sugar and is tuned by the CD and FG loops of the V-set Ig fold (**Fig. 4, 5, S5–S11**). One advantage of this architecture is that the likely flexible trisaccharides can productively interact with the binding pocket via multiple approaches, i.e. binding the sialic acid first or by binding the reducing terminus of the glycan first. The concept of a binding site with multiple independent recognition regions can be extrapolated to adhesins that recognize larger sialoglycans. For example, the Siglec-like adhesin SrpA may biologically recognize a hexasaccharide (17) but can bind to partial ligands, albeit with low affinity (16, 18, 20) (**Fig. 2C**).

Mutagenesis (**Fig. S8**), chimeragenesis (**Fig. 4, S9, S10**), and computer simulations (**Fig. 3, S7**) all suggest that flexibility of these loops controls the breadth of the binding spectrum via a conformational selection mechanism. Binding promiscuity correlates with the identity of the EF loop (**Fig. S9**), which suggests a mechanism where the EF loop adjusts ligand orientation. The variable region of the ligand can then approach the CD and FG selectivity loops at different angles in order to optimize interactions with the myriad of positions of hydrogen-bonding donors and acceptors that decorate the diverse glycans recognized. If the ligand binds to the CD and FG loops first, the order of events would be reversed, but the mechanism unchanged.

Chimeragenesis and mutagenesis also indicate that the CD and FG loops are particularly important in determining the preferred ligand (**Fig. 4, 5, S9 – S11**). The use of loops to control selectivity has previously been observed in other sialoglycan-binding systems. For example, the mammalian Siglec proteins are built upon a V-set Ig-fold but are not detectably related in sequence to the SRR adhesins (20, 21, 31, 32). In Siglec-7, the CC’ loop (33) controls sialoglycan selectivity. From an evolutionary standpoint, having a mutable loop control selectivity makes particular sense for oral bacteria because it allows facile alteration of ligand preference in response to a changing environment. Indeed, mutation of loops is unlikely to impact protein stability.

The more promiscuous Hsa-like adhesins appeared to be particularly amenable to engineering (**Fig. 4, 5, S11**) and mutants exhibited binding increases of 2- to 3-orders of magnitude for non-native ligands. These increases exceed those reported for dedicated engineering studies (34-42), where the maximum enhancement in binding to a non-native glycan is ∼20-fold (34-39) but selectivity is often achieved via a decrease in affinity to non-desired ligands in a promiscuous starting lectin (40-42). One intriguing interpretation of the unusually facile engineering of these Siglec-like adhesins is that their biological role necessitates adjusting to changes in host environment. An easily mutable adhesin may confer a survival advantage by allowing a bacterial strain to adapt to changes in the glycan modifications on salivary MUC7, adapt to binding to distinct receptors in a new anatomical location, or even to adapt to a new host. One impact of adaptation to different preferred receptors could be the ability of these bacteria to convert from commensals to pathogens (26).

An exciting outcome is our engineering of adhesins selective for sTa (**Fig. 4E**) and 6S-sLe^X^ (**Fig. 5A, 5B**) on human proteins (**Fig. 6**). Adhesins with novel sialoglycan selectivity have multiple potential applications. The inherent challenges associated with characterizing *O*-glycans leave many biological questions arising from knowledge of sialoglycan distribution under-addressed. One strategy for mapping the glycome has been to repurpose naturally-occurring glycan-binding proteins as probes (43). Engineered probes could expand the range of detectable glycans. A second application is in detecting altered glycosylation in disease. Overexpression of sialoglycans is a biomarker for many types of cancers and commonly associated with poor prognosis (44-49). Robust antibodies to many sialoglycans, in particular sialyl-Thompson-nouvelle antigen (sTn) have proven a challenge to develop (50). One could envision highly-selective lectins being used for detection of sialoglycans via lectin-based microarrays (43) or ELISAs. These may also be used in histological mapping or affinity purification of specific protein glycoforms. Additional work could develop probes selective for other Siaα2-3Gal-linked sialoglycans or for sialoglycans with other linkages and is a future goal.

## Methods

Detailed procedures for protein expression, purification, crystallization, structure determination, computational analyses, and binding assays are described in the SI.

### Accession numbers

Atomic coordinates and structure factors have been deposited into the RCSB Protein Data Bank and raw data have been deposited into SBGrid with accession codes listed in **Tables S1 and S2**.

## Acknowledgements

We thank S Bordenstein for assistance with phylogenetic analysis and B Bachmann, R Woods, and O Grant for helpful discussions. This work was supported by the Department of Veterans Affairs, the National Institutes of Health (AI41513 and U01CA221244 to PS, HD065122 to XC, AI106987 to TMI/PMS, and DE019807 to TMI), and the American Heart Association (14GRNT20390021 to TMI; 17SDG33660424 to BAB). AH was supported by the Vanderbilt International Scholars Program, and MAC was supported by a National Science Foundation Individual pre-doctoral fellowship (DGE-1445197). Use of the Stanford Synchrotron Radiation Lightsource, SLAC National Accelerator Laboratory, is supported by the U.S. Department of Energy, Office of Science, Office of Basic Energy Sciences under Contract No. DE-AC02-76SF00515. The SSRL Structural Molecular Biology Program is supported by the DOE Office of Biological and Environmental Research, and by the National Institutes of Health, National Institute of General Medical Sciences (including P41GM103393). The Advanced Photon Source, a User Facility operated for the U.S. DOE Office of Science, was supported under Contract DE-AC02-06CH11357. LS-CAT Sector 21 is supported by the Michigan Economic Development Corporation and the Michigan Technology Tri-Corridor (085P1000817).

## Author contributions

TMI and BAB designed research, BAB, LVL, IY, KL, AH, RA and ZW performed research, BAB, MAC, KL, IY, KF and TMI analyzed data, HY and XC contributed reagents/analytical tools. JB, JCS, PMS and TMI conceived and guided the study. TMI wrote the manuscript. All authors approved the final version of the manuscript.

## Supporting Information

### Materials and Methods

#### Sequence analysis

Sequences of the tandem Siglec and Unique domains were resected from select adhesins and were aligned using the MUSCLE (1) subroutine in Geneious Pro 11.1.4 (2). The JTT-G model of evolution was selected using the ProtTest server (3), and the phylogenetic tree was built using the MrBayes (4) subroutine in Geneious Pro 11.1.4 (2). A distantly-related adhesin from *S. mitis* strain SF100 (5) was used to root the tree.

#### Cloning, expression, and purification for crystallization

DNA encoding the adjacent Siglec and Unique domains of GspB, SK150, NCTC10712, or SK678 or the Siglec domain of GspB were cloned into the pBG101 vector (Vanderbilt University), which encodes an N-terminal His_6_-GST tag that is cleavable using 3C protease. Hsa_Siglec+Unique_ was cloned into the pSV278 vector (Vanderbilt University), which encodes a His_6_-maltose binding protein (MBP) tag at the N-terminus followed by a thrombin cleavage site. Proteins were expressed in *E. coli* BL21 (*DE3*) in Terrific Broth medium (for GspB proteins and Hsa_Siglec+Unique_) or LB (for SK150_Siglec+Unique_, NCTC_Siglec+Unique_ and SK678_Siglec+Unique_) with 50 µg/ml kanamycin at 37°C. When the OD_600_ reached 0.6-1.4, expression was induced with 0.5-1 mM IPTG at 24°C for 3-7 hrs. Cells were harvested by centrifugation at 5,000 × *g* for 15 min, optionally washed with 0.1 M Tris-HCl, pH 7.5, and stored at –20 °C before purification.

Frozen cells were resuspended in homogenization buffer (20-50 mM Tris-HCl, pH 7.5, 150-200 mM NaCl, 1mM EDTA, 1 mM PMSF, 2 µg/ml Leupeptin, 2 µg/ml Pepstatin) then disrupted by sonication. Lysate was clarified by centrifugation at 38500 × *g* for 35-60 min and passed through a 0.45 µm filter. Purification was performed at 4 °C. His_6_-GspB-fusion proteins were purified using a Glutathione Sepharose 4B column and were eluted with 30 mM GSH in 50 mM Tris-HCl, pH 8.0. His_6_-SK150_Siglec+Unique_/NCTC10712_Siglec+Unique_/SK678_Siglec+Unique_ proteins were purified using Ni^2+^ affinity chromatography and eluted with 20 mM Tris-HCl, 150 mM NaCl, 250 mM imidazole, pH 7.6. His_6_-MBP-Hsa_Siglec+Unique_ was purified with an MBP-Trap column and eluted in 10 mM maltose. Eluted proteins were concentrated in a 10kD MW cut-off concentrator and exchanged into either PreScission cleavage buffer (GspB_Sigelc_, GspB_Siglec+Unique_, SK150_Siglec+Unique_, NCTC10712_Siglec+Unique_, or SK678_Siglec+Unique_; 50 mM Tris-HCl, pH 7.6, 150 mM NaCl, 1mM DTT) or thrombin cleavage buffer (Hsa_Siglec+Unique_; 20 mM Tris-HCl pH 7.5 and 200 mM NaCl). Affinity tags were cleaved with 1 U of appropriate protease (thrombin or 3C) per mg of protein overnight at 4 °C. For the SK150_Siglec+Unique_, NCTC10712_Siglec+Unique_, and SK678_Siglec+Unique_, the affinity tag has a similar molecular weight as the target protein; in these cases, the cleaved sample was passed through a Ni-column to remove the His_6_-GST tag. For GspB domains, adhesin was separated from the affinity tag by passing the cleavage reaction over the second Glutathione Sepharose 4B column in PreScission Buffer. Protein aggregates were removed from GspB domains using a Superose-12 column in 50 mM Tris-HCl pH 7.6 and 150 mM NaCl. For the remaining proteins, aggregates were removed using a Superdex 200 increase 10/30 GL column equilibrated in 20 mM Tris-HCl pH 7.6 (NCTC_Siglec+Unique_, SK150_Siglec+Unique_, SK678_Siglec+Unique_) or in 20 mM Tris-HCl pH 7.5 and 200 mM NaCl (Hsa_BR_). After purification, all proteins were >95% pure as assessed by SDS-PAGE and were stored at −80 °C.

#### Crystallization, data collection, and structure determination

All crystallization reactions were performed at room temperature (∼23 °C). Unless otherwise noted, diffraction data were collected at - 180 °C, processed using HKL200 (6), and structures were determined by molecular replacement using the Phaser (7) subroutine of Phenix (8) and the search model indicated. Riding hydrogens were included at resolutions better than 1.4 Å. X-ray sources and data collection statistics are found in **Supporting Tables 1 & 2**.

##### GspB

GspB domains were crystallized by the sitting drop vapor diffusion method by equilibrating 1 µL protein and 1 µL reservoir solution over 50 µL of a reservoir solution. Purified GspB_Siglec+Unique_ was concentrated to 9 mg/ml in 20 mM Tris-HCl, pH 7.6 and crystallized using a reservoir containing 0.2 M (NH_4_)_2_SO_4_, 25% polyethylene glycol (PEG) 3350. Crystals were flash cooled by plunging into liquid nitrogen without the addition of cryo protectant. Purified GspB_Siglec_ was concentrated to 22.8 mg/ml in 20 mM Tris-HCl, pH 7.2. Crystals in space group P2_1_2_1_2 were grown with a reservoir solution containing 0.2 M MgCl_2_, 0.1 M Tris-HCl, pH 8.5, 30% w/v PEG 4000; crystals in space group R32 were grown with a reservoir containing 4.0 M HCOONa. GspB_Siglec_ was cocrystallized with sTa using reservoir conditions associated with the P2_1_2_1_2 space group and 1 µL of protein-ligand complex (20.5 mg/ml GspB_Siglec_, 10 mM sTa, 18 mM Tris-HCl, pH 7.2). Structures were determined using the appropriate domain(s) of GspB (PDB entry 3QC5 (9)) resected from the three-domain structure.

##### SK150

Purified SK150_Siglec+Unique_ was concentrated to 3.5 mg/ml in 20 mM Tris-HCl, pH 7.6. Crystals were grown by the hanging drop vapor diffusion method by mixing 1 µL protein and 1 µL reservoir solution (0.2 M ammonium sulfate, 25% PEG 4000, 15% ethanol, and 0.1M Bis-tris, pH 7.0) and equilibrating over the reservoir solution. Diffraction data were collected at room temperature (∼23 °C) and were processed using the PROTEUM suite. The structure was determined using the Siglec and Unique domains of GspB (PDB entry 3QC5 (9)) as the search model.

##### Hsa

Crystals of Hsa_Siglec+Unique_ (21.6 mg/ml in 20 mM Tris-HCl, pH 7.2) grew by sitting drop vapor diffusion by equilibrating 1 µL protein and 2 µL reservoir solution over 50 µL of reservoir solution (0.1 M Succinate/Phosphate/Glycine pH 10.0 and 25% PEG 3350). Co-crystals of Hsa_Siglec+Unique_ with sTa were prepared by soaking fully formed crystals in reservoir solution supplemented with 5 mM sTa for 20 hr. Crystals did not require cryoprotection beyond the reservoir solution. The structure of unliganded Hsa_BR_ was determined using *S. sanguinis* SrpA_Siglec+Unique_ (PDB entry 5EQ2 (10)) as the search model. The structure of sTa-bound Hsa_BR_ was determined by rigid body refinement of unliganded Hsa_Siglec+Unique_ in Phenix (8).

##### NCTC10712

Crystals of NCTC10712_Siglec+Unique_ (3.5 mg/ml in 20 mM Tris-HCl pH 7.5) grew via the hanging drop vapor diffusion method using reservoir containing 0.1 M Tris-HCl pH 7.5 and 32% w/v PEG 4000. Crystal quality was improved by microseeding (Hampton Seed Bead kit) using 0.3 µL of seed, 1.2 µL protein (3.5 mg/ml), and 1.5 µL modified reservoir solution (0.1 M Tris-HCl pH 7.5 and 28% w/v PEG 4000). Crystals were cryoprotected in using a solution containing 50% of the reservoir and 50% glycerol, then cryocooled by plunging in liquid nitrogen. Data were processed using XDS (11). The structure was determined Hsa_Siglec+Unique_ as the search model.

##### SK678

Crystals of SK678_Siglec+Unique_ (7 mg/ml in 20 mM Tris-HCl pH 7.6) were grown via the hanging drop vapor diffusion method by equilibrating 1 µL of SK678_Siglec+Unique_ and 1 µL reservoir solution over the reservoir solution (0.1M Bicine pH 7.6 and 25% PEG 6,000, 0.005M hexamine cobalt(II) chloride). Crystals were cryoprotected in artificial reservoir solution containing 15% glycerol, and 15% ethylene glycol, then cryo cooled by plunging into liquid nitrogen. Diffraction data were processed using XDS (11). The structure was determined using NCTC10712_BR_ as the search model.

#### Crystallographic refinement, and analysis

All models were improved with iterative rounds of model building in Coot (12) and refinement in Phenix (8). In all structures of GspB subdomains, the unliganded structure of Hsa_BR_, and the structure of NCTC_BR,_ electron density for hydrogens was observed in later rounds of refinement and riding hydrogens were included in the final model, which reduced the R_free_ by over 1% in each case. Bound cations were assigned as either Na^+^, Mg^2+^, or Ca^2+^ depending upon the abundance of these ions in either the purification or the crystallization conditions, and the previous observation that cations bound to this site are readily exchanged with cations in the buffer (9). The final models are associated with the statistics listed in **Supporting Tables 1** and **2**. When Ramachandran outliers are associated with the models, these are unambiguously defined by clear electron density.

For sTa-bound Hsa_Siglec+Unique_ and GspB_Siglec_, the crystals were isomorphous with unliganded crystals. Accordingly, R_free_ reflections were selected as identical. In both cases, unambiguous electron density for all three sugars of sTa was apparent in the initial maps. Ligand occupancies were held at 1.0 during refinement.

#### Sialoglycan binding assays

DNA encoding wild-type and variant adhesins were cloned into pGEX-3X. Chimeras were designed using an overlay of the coordinates from each adhesin crystal structure. DNA encoding adhesin chimeras were cloned into pGEX-3X. SK678-Hsa chimeras had the Siglec and Unique domains of SK678 and the loops from Hsa. GspB-SK150 chimeras had the Siglec and Unique domains of GspB with selectivity loops of SK150.

The pGEX vectors encode an N-terminal glutathione *S*-transferase (GST) affinity tag, which was used for purification. Individual GST-Siglec+Unique fusions were expressed and purified using glutathione-sepharose, and the binding of biotinylated glycans to immobilized GST-binding regions was performed as described previously (5).

#### Far Western and lectin blotting of human plasma proteins

Far-western blotting of human plasma proteins using the indicated GST-binding regions (15 nM) as probes was performed as described (13).

#### Interdomain angle calculations

The torsion angle between Siglec and Unique domains for each system (GspB, SK150, Hsa, SK678, NCTC10712, SrpA) were defined as the angle between the planes formed between center of mass (COM) of Siglec and Residue 1 (R1) and COM of Unique and Residue 2 (R2). The two residues (R1 & R2) were chosen based on crystal structure alignment and are listed in **Table S4**. Missing residues of SK150_Siglec+Unique_ were modeled using GspB_Siglec+Unique_ as a template (PDB entry 3QC5, (9)).

#### Molecular dynamics (MD) simulations and analyses

For MD simulations, each system (GspB or Hsa) was solvated in a 10 Å octahedral box of TIP3P (14) water. The Amber16 ff14SB (15) force field was used for the protein. In the first step of the MD simulation, the backbone and side chains of the protein was restrained using 500 kcal mol^-1^ Å^-2^ harmonic potentials while the system was energy minimized for 500 steps of steepest descent (16). This step was followed by 500 steps with the conjugate gradient method (17). In a second minimization step, restraints on the protein were removed and 1000 steps of steepest descent minimization were performed followed by 1500 steps of conjugate gradient. The system was then subjected to MD and heated to 300 K with the backbone and side chains of the protein restrained using 10 kcal mol^-1^ Å^-2^ harmonic potentials for 1000 steps. The restraints were released and 1000 MD steps were performed. The SHAKE(18) algorithm was used to constrain all bonds involving hydrogen in the simulations. MD runs (200 ns) were performed at 300 K in the NPT ensemble and a 2 fs time step. The probability distribution analyses and RMSF calculations were performed on 200 ns of 3 independent runs for each system. All analyses were performed using the cpptraj and pytraj (19) python modules of AMBER16.

---

**Table S1.** Crystallographic data collection and refinement statistics for Hsa-like adhesins. Values in parentheses are for the highest resolution shell. Raw data are deposited with SBGrid and can be accessed at: data.sbgrid.org/dataset/DATAID. The main chain of Hsa_Siglec+Unique_ displays two alternative conformations between residues 378-384 that is accompanied by a *cis-trans* isomerization of the non-proline peptide bond between Hsa^E381^ and Hsa^S382^ and results in disallowed bond angles for Hsa^S383^. In other isoforms of Hsa, the equivalent residue is a proline. Current technologies do not allow main chain alternative conformations to be refined within the same model. One position is modeled in the unliganded Hsa_Siglec+Unique_ structure and one conformation is modeled in the sTa-bound Hsa_Siglec+Unique_ structure; however, electron density for both conformations is clearly visible in both structures. *The Ramachandran angles identified as outliers (Hsa^S253^, Hsa^L363^, NCTC10712^S253^, NCTC10712^L361^) are equivalent in the homologs and are associated with clear electron density.

|  | **Hsa<sup>Siglec+Unique</sup>** | **Hsa<sup>Siglec+Unique</sup> +**<br>**sTa** | **NCTC10712<sup>Siglec+Unique</sup>** | **SK678<sup>Siglec+Unique</sup>** |
| --- | --- | --- | --- | --- |
| PDB entry | 6EFC | 6EFD | 6EFF | 6EFI |
| DATAID | 328 | 329 | 509 | 510 |
| Resolution | 1.4 Å | 1.85 Å | 1.6 Å | 1.7 Å |
| *Data collection* |  |  |  |  |
| Beamline | APS 21-ID-F | APS 21-ID-G | APS 21-ID-F | SSRL 9-2 |
| Wavelength | 0.978 Å | 0.978 Å | 0.978 Å | 0.979 Å |
| Space group | P2<sub>1</sub>2<sub>1</sub>2<sub>1</sub> | P2<sub>1</sub>2<sub>1</sub>2<sub>1</sub> | P1 | P2<sub>1</sub> |
| Unit cell | a=46.6 Å<br>b=58.1 Å<br>c=76.0 Å | a=46.7 Å<br>b=58.0 Å<br>c=76.1 Å | a=39.8 Å<br>b=48.9 Å<br>c=99.8 Å<br>$\alpha=101.8^\circ$<br>$\beta=91.4^\circ$<br>$\gamma=89.9^\circ$ | a=59.6 Å<br>b=59.58 Å<br>c=61.8 Å<br>$\beta=100.7^\circ$ |
| R<sub>sym</sub> | 0.084 (0.650) | 0.107 (0.638) | 0.075 (0.730) | 0.099 (0.530) |
| R<sub>pim</sub> | 0.024 (0.281) | 0.037 (0.218) | 0.047 (0.479) | 0.040 (0.213) |
| I/$\sigma$ | 49.7 (2.3) | 31.7 (2.9) | 22.9 (1.9) | 15.0 (4.4) |
| Completeness (%) | 93.3% (60.9%) | 98.8 % (89.5%) | 92.4% (70.9%) | 97.7% (97.3%) |
| Redundancy | 12.6 (5.6) | 9.5 (8.7) | 3.6 (3.4) | 7.0 (7.1) |
| CC<sub>1/2</sub> | 0.837 | 0.940 | 0.648 | 0.998 |
| *Refinement* |  |  |  |  |
| R<sub>cryst</sub> | 0.146<sup>&</sup> | 0.196<sup>&</sup> | 0.180 | 0.177 |
| R<sub>free</sub> | 0.179 | 0.217 | 0.207 | 0.210 |
| No. Mol per ASU | 1 | 1 | 4 | 2 |
| RMS deviation |  |  |  |  |
| bond lengths | 0.01 Å | 0.01 Å | 0.01 Å | 0.01 Å |
| bond angles | 1.6° | 0.9° | 0.9° | 0.7° |
| Ramachandran |  |  |  |  |
| favored | 97.0% | 97.1% | 96.8% | 99.0% |
| allowed | 2.5% | 2.9% | 3.1% | 1.0% |
| outliers | 0.5%<sup>*</sup> | 0.0%<sup>*</sup> | 0.1%<sup>*</sup> | 0.0% |

**Table S2.** Crystallographic data collection and refinement statistics for GspB-like adhesins. Values in parentheses are for the highest resolution shell. Raw data are deposited with SBGrid and can be accessed at: data.sbgrid.org/dataset/DATAID

|  | **GspB<sub>Siglec</sub>** | **GspB<sub>Siglec</sub> +**<br>**sTa** | **GspB<sub>Siglec</sub>**<br>**Form 2** | **GspB<sub>Siglec+Unique</sub>** | **SK150<sub>Siglec+Unique</sub>** |
| --- | --- | --- | --- | --- | --- |
| PDB entry | 6EF7 | 5IUC | 6EF9 | 6EFA | 6EFB |
| DATAID |  | 507 | 601 | 604 | 508 |
| Resolution | 1.03 Å | 1.25 Å | 1.3 Å | 1.6 Å | 1.90 Å |
| *<u>Data collection</u>* |  |  |  |  |  |
| Beamline | 21-ID-F | 21-ID-F | 21-ID-G | 21-ID-F | Bruker X8R |
| Wavelength | 0.979 Å | 0.979 Å | 0.979 Å | 0.979 Å | 1.542 Å |
| Space group | P2<sub>1</sub>2<sub>1</sub>2 | P2<sub>1</sub>2<sub>1</sub>2 | R32 | P2<sub>1</sub>2<sub>1</sub>2<sub>1</sub> | P2<sub>1</sub> |
| Unit cell | a= 33.9 Å | a=67.7 Å | a=b=92.1 Å | a=33.0 Å | a=24.3 Å |
|  | b= 46.2 Å | b=66.6 Å |  | b=47.6 Å | b=62.6 Å |
|  | c= 73.0 Å | c=55.9 Å | c=143.9 Å | c=136.2 Å | c=62.9 Å |
|  |  |  |  |  | β=98.6° |
| R<sub>sym</sub> | 0.049 (0.430) | 0.066 (0.406) | 0.061 (0.771) | 0.057 (0.610) | 0.139 (0.538) |
| R<sub>pim</sub> | 0.024 (0.233) | 0.018 (0.111) | 0.017 (0.302) | 0.019 (0.213) | 0.044 (0.295) |
| I/σ | 25.5 (4.4) | 35.6 (7.9) | 59.3 (2.8) | 43.8 (3.3) | 9.3 (1.9) |
| Completeness (%) | 95.4% (90.6%) | 95.7% (90.5%) | 99.9% (98.0%) | 88.9% (48.6%) | 97.3% (91.6%) |
| Redundancy | 9.3 (8.2) | 14.9 (14.3) | 13.8 (7.1) | 9.2 (7.7) | 9.0 (3.6) |
| CC<sub>1/2</sub> | 0.911 | 0.964 | 0.941 | 0.975 | 0.996 |
| *<u>Refinement</u>* |  |  |  |  |  |
| R<sub>cryst</sub> | 0.125 | 0.156 | 0.131 | 0.166 | 0.172 |
| R<sub>free</sub> | 0.141 | 0.178 | 0.144 | 0.209 | 0.188 |
| No. Mol per ASU | 1 | 2 | 2 | 1 | 1 |
| RMS deviation |  |  |  |  |  |
| bond lengths | 0.01 Å | 0.01 Å | 0.01 Å | 0.02 Å | 0.01 Å |
| bond angles | 1.3° | 1.5° | 1.1° | 1.6° | 0.7° |
| Ramachandran |  |  |  |  |  |
| favored | 100% | 99.2% | 97.0% | 98.0% | 99.0% |
| allowed | 0% | 0.8% | 2.2% | 1.5% | 1.0% |
| outliers* | 0% | 0% | 0.8% | 0.5% | 0.0% |

**Table S3.** Summary of binding preferences of wild-type and variant SRR adhesins. sTa, sialyl-T antigen, 3’sLn, 3’-sialyl-*N*-acetyllactosamine, sLe^C^, sialyl-Lewis^C^, sLe^X^, sialyl-Lewis^X^, 6S-sLe^X^, 6-*O*-sulfo-sialyl Lewis^X^, nd= not determined

|  | sTa | sLe<sup>C</sup> | 3'sLn | sLe<sup>X</sup> | 6S-sLe<sup>X</sup> |
| --- | --- | --- | --- | --- | --- |
| **Hsa**<sub>Siglec+Unique</sub> | +++ | ++ | ++ | + | + |
| **S253G** | +++ | nd | ++ | nd | nd |
| **L363G** | +++ | nd | ++ | nd | nd |
| **N333P** | ++ | nd | - | nd | nd |
| **G287A/G288P** | +++ | nd | ++ | nd | nd |
| **E286R** | +++ | +++ | +++ | +++ | +++ |
| **D356Q** | ++ | ++ | +++ | ++ | +++ |
| **D356R** | + | ++ | +++ | + | +++ |
| **NCTC10712**<sub>Siglec+Unique</sub> | + | + | +++ | ++ | +++ |
| **E285R** | - | - | + | - | +++ |
| **Q354D** | + | - | + | - | - |
| **+ Hsa CD loop** | + | + | + | + | +++ |
| **+ Hsa EF loop** | - | + | +++ | +++ | +++ |
| **+ Hsa FG loop** | ++ | + | ++ | + | + |
| **+ all Hsa loops** | +++ | ++ | + | - | - |
| **SK678**<sub>Siglec+Unique</sub> | - | - | ++ | + | ++ |
| **E302R** | - | - | + | - | +++ |
| **Q371D** | - | - | + | - | - |
| **+ Hsa CD loop** | - | - | - | - | - |
| **+ Hsa EF loop** | - | - | ++ | + | ++ |
| **+ Hsa FG loop** | - | - | + | - | - |
| **+ all Hsa loops** | ++ | + | + | - | - |
| **GspB**<sub>Siglec+Unique</sub> | +++ | - | - | - | nd |
| **L442Y/Y443N** | ++ | + | ++ | nd | nd |
| **+ SK150 CD loop** | - | nd | - | nd | nd |
| **+ SK150 EF loop** | +++ | nd | - | nd | nd |
| **+ SK150 FG loop** | - | nd | - | nd | nd |
| **+ all SK150 loops** | - | nd | - | nd | nd |
| **SK150**<sub>Siglec+Unique</sub> | ++ | + | + | - | - |
| **Y300L/N301Y** | - | nd | - | nd | nd |

**Table S4.** Residues used for interdomain angle calculations.

| **Protein** | **Residues (R1 & R2)** |
| --- | --- |
| **GspB** | **Glu 417, Asp 557** |
| **SK150** | **Glu 275, Asn 416** |
| **Hsa** | **Glu 262, Asn 411** |
| **SK678** | **Glu 274, Gln 422** |
| **NCTC10712** | **Glu 261, Ser 409** |
| **SrpA** | **Glu 271, Gln 409** |

**Figure S1.**
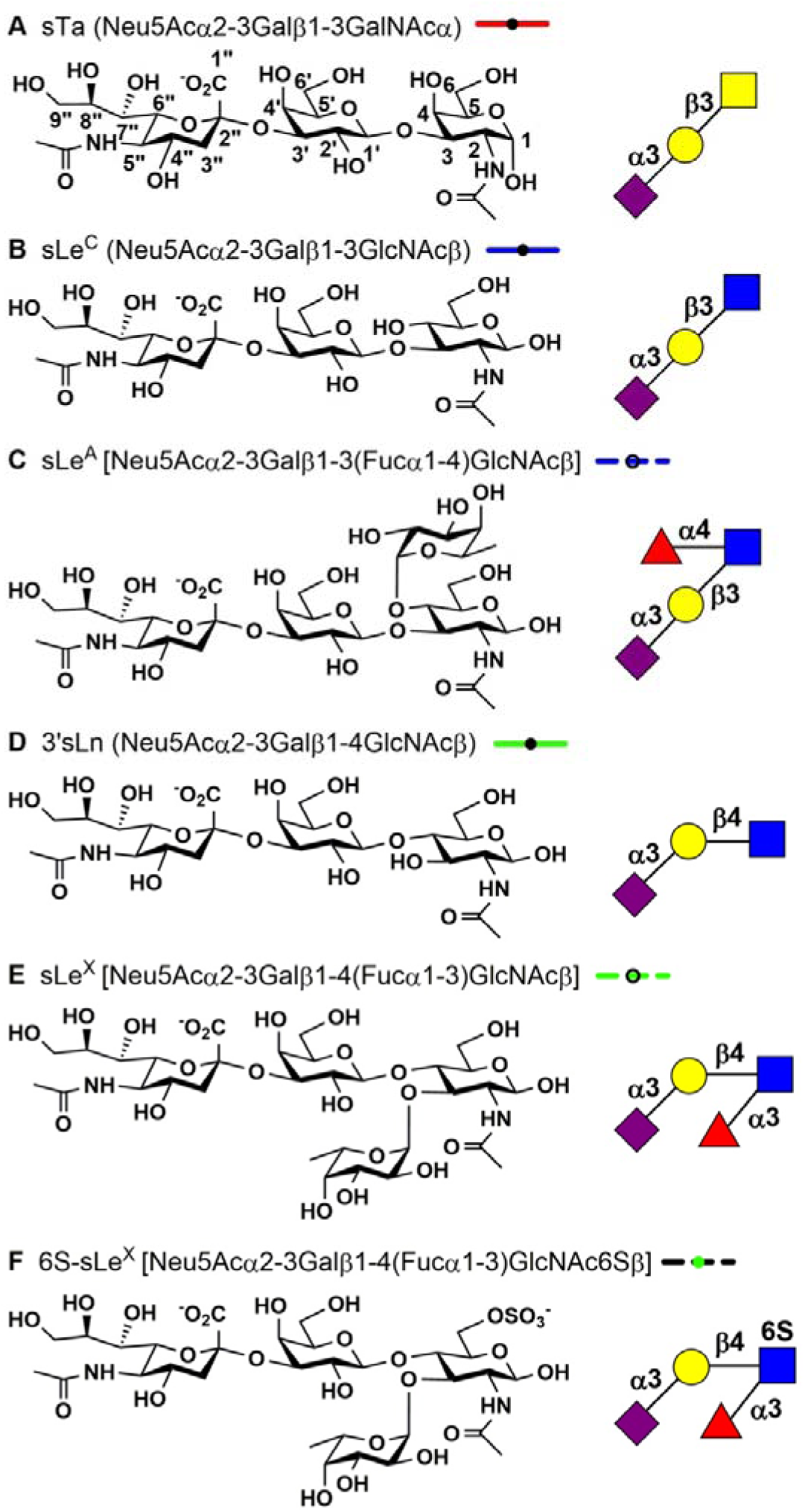
Sialoglycans used in this study. The chemical structure of each indicated sialoglycan is shown of the left with the symbolic representation shown on the right. The line style used for all dose response curves is shown to the right of each name.

**Figure S2.**
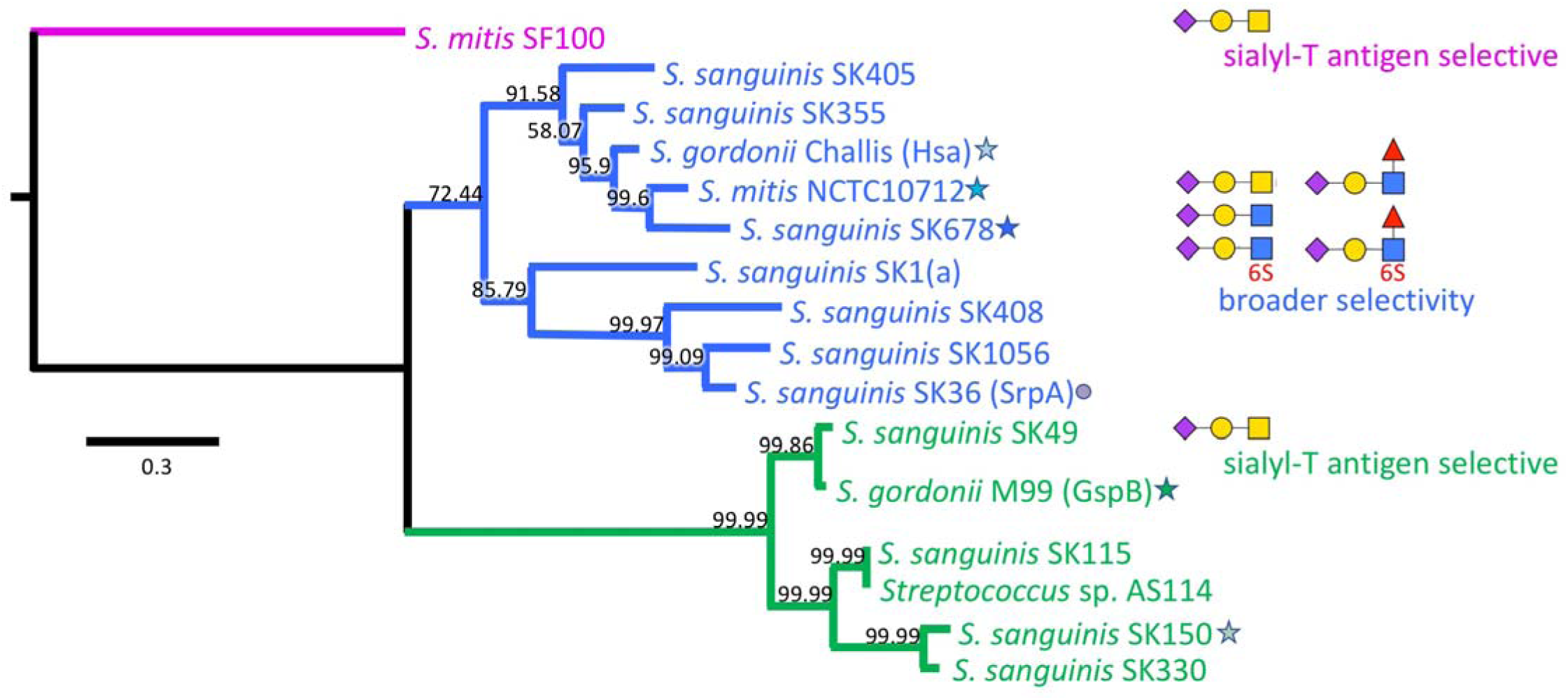
Bacterial Siglec-like SRR adhesins. Phylogenetic analysis of the tandem Siglec and Unique domains of select bacterial SRR adhesins reveals three distinct subgroups. Characterized Hsa-like adhesins (*blue*) bind to two or more of the indicated sialoglycans; the four characterized GspB-like adhesins (*green*) have narrow selectivity for sialyl-T antigen. The tree is rooted using the distantly-related *S. mitis* SF100 adhesin (*magenta*). Adhesins investigated here are highlighted with a star, and figure panels comparing properties of these adhesins follow this coloring. The structure and ligand binding properties of SrpA, highlighted with a circle, have previously been reported(10, 20), and SrpA is used as a comparator in this report.

**Figure S3.**
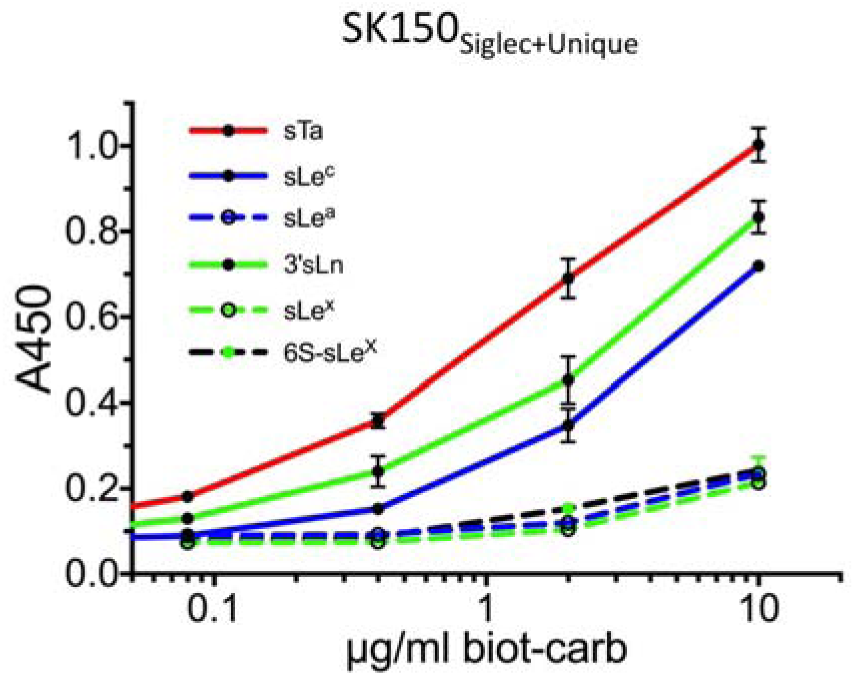
Binding of select trisaccharides to GST-SK150_Siglec+Unique_. GST-SK150_Siglec+Unique_ (500 nM) was immobilized in 96-well plates, and biotinylated sialoglycan ligands were added at the indicated concentrations. Binding is reported as the mean ± standard deviation, with n = 3.

**Figure S4.**
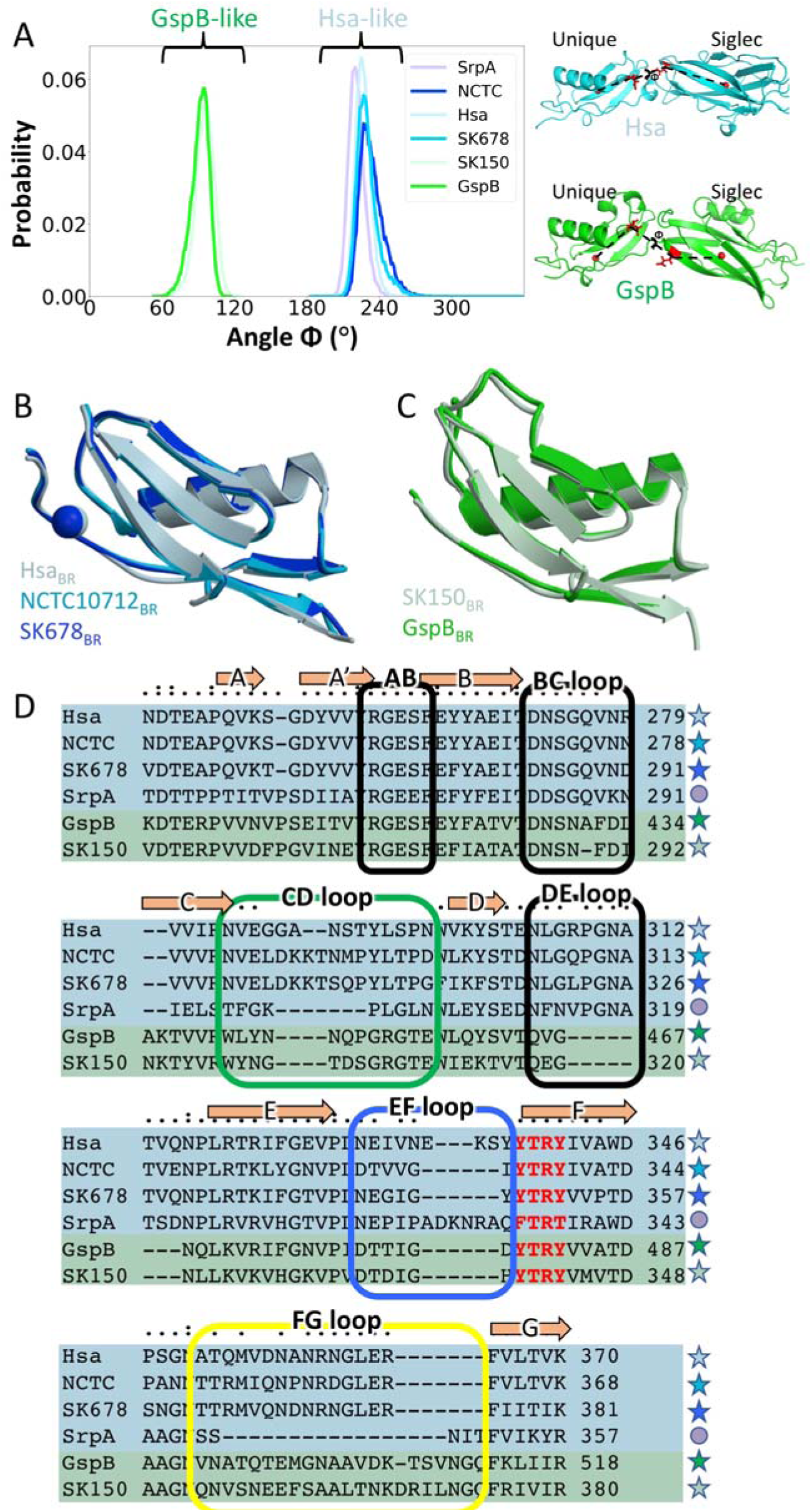
Comparisons of bacterial SRR adhesins. A. Probability distributions of the torsion angle (Φ) between the Siglec and Unique domains of GspB, SK150, Hsa, NCTC10712, SK678, SrpA (left) as calculated from MD simulations. Crystal structures showing the Φ angle for both GspB-like and Hsa-like proteins (right). The interdomain torsion angles in the corresponding crystal structures are as follows: GspB: ∼100°; SK150: ∼100°; NCTC10712: ∼228°; Hsa: ∼230°; SrpA: ∼216°; SK678: ∼240°. B, C. Overlay of the Unique domains of: B. Hsa-like adhesins and C. GspB-like adhesins. The view is rotated as compared to **Fig. 2** in order to highlight the structural similarity between the branches of the phylogenetic tree. D. Sequence alignment of the Siglec domain of SRR adhesins. Hsa-like adhesins are highlighted with a blue background and GspB-liked adhesins are highlighted with a green background. Strands conserved in the V-set Ig fold are indicated, and residues of the interstrand loops are highlighted with boxes.

**Figure S5.**
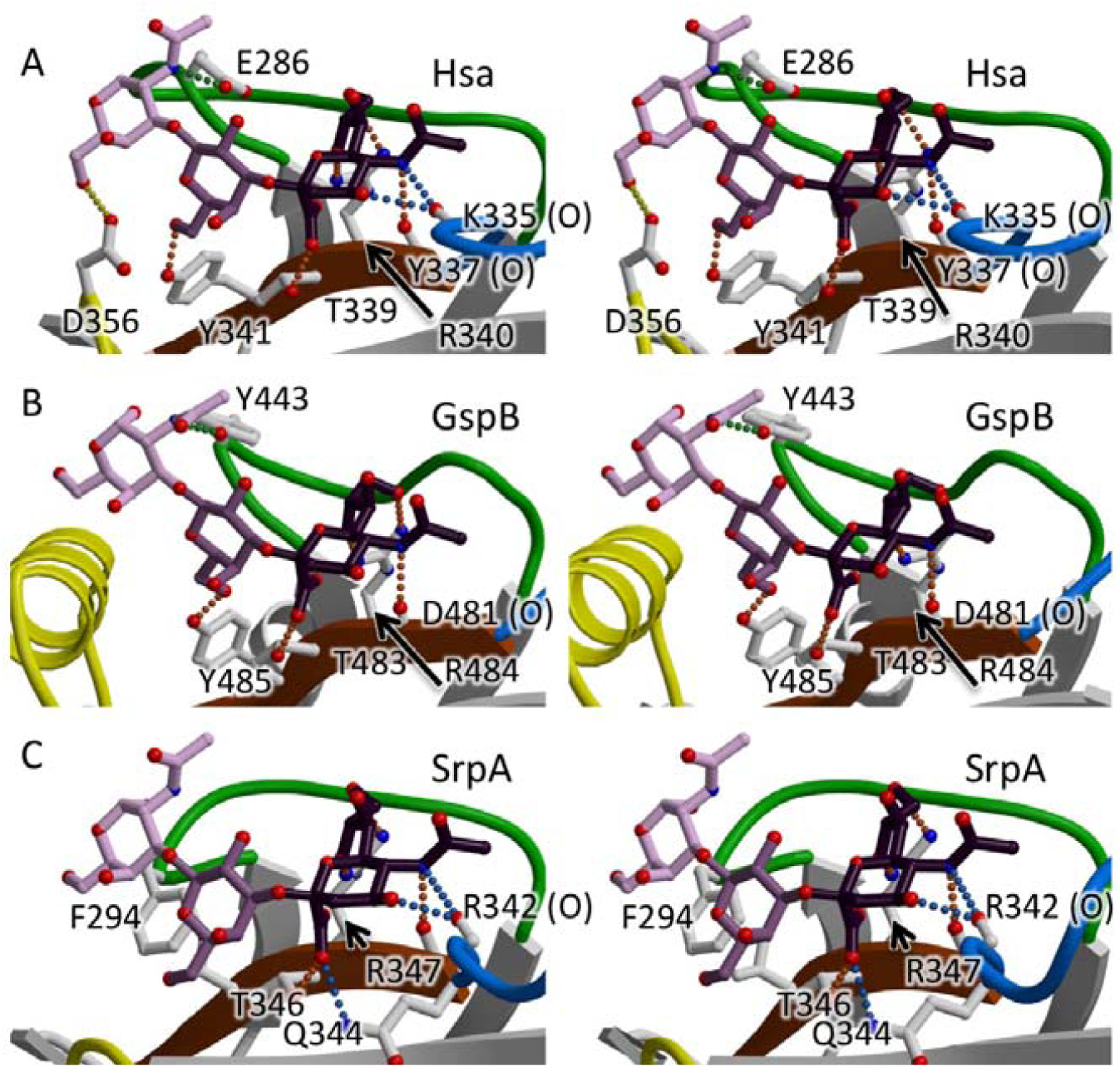
Stereoviews of the contacts between bacterial SRR adhesins and the sTa ligand. A. Hsa_Siglec+Unique_ bound to sTa. B. GspB_Siglec_ bound to sTa. C. SrpA_Siglec+Unique_ bound to sTa (PDB entry 5IJ3 (10)). Residues of each respective adhesin within hydrogen-bonding distance of sTa are labeled. Color scheme follows that of Fig. 3 with the CD loop in *green*, the EF loop in *blue*, and the FG loop in *yellow*.

**Figure S6.**
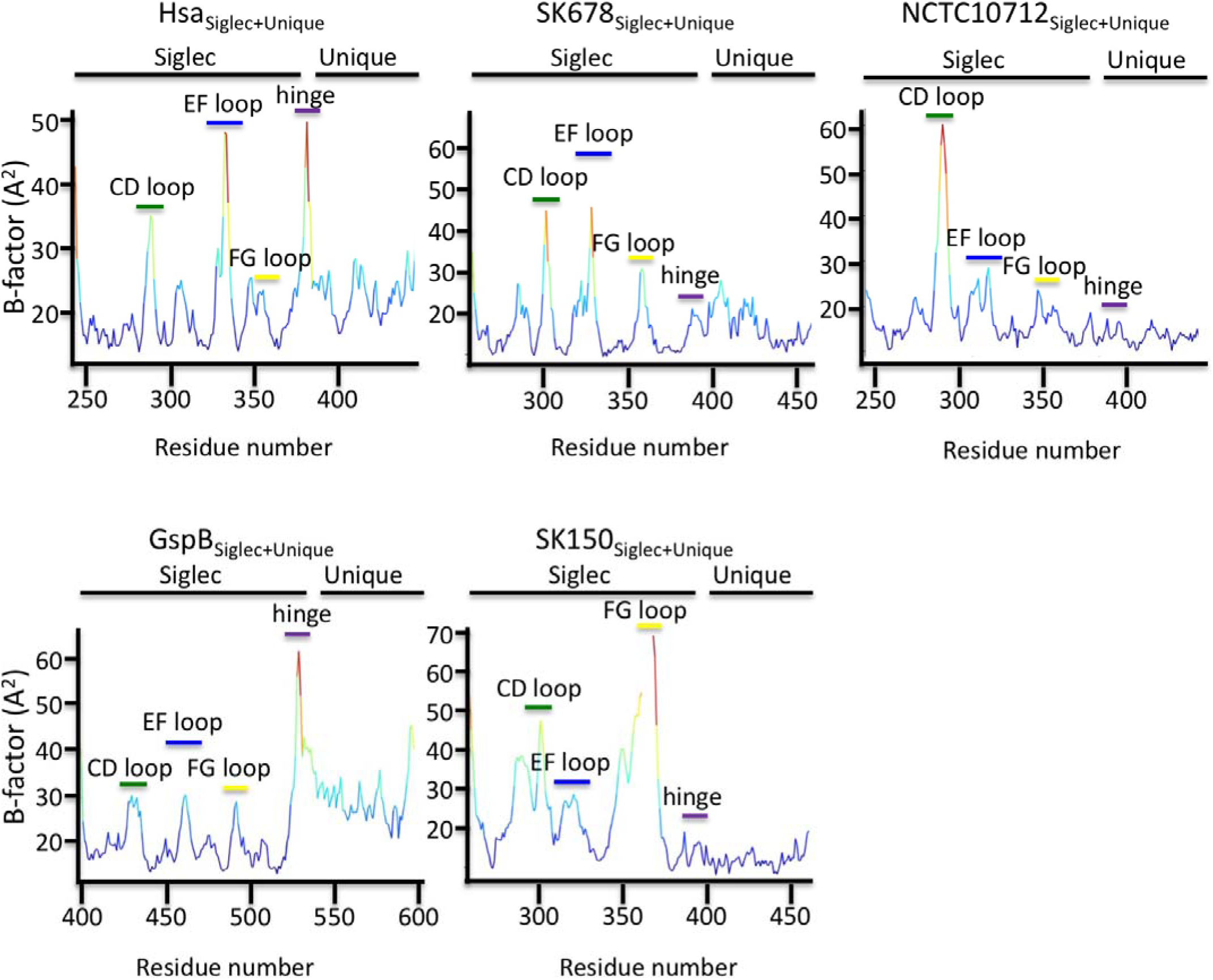
Temperature factor analysis of adhesins. For each graph, the residue number is on the x-axis, and the crystallographic temperature factor (B-factor) is on the y-axis. Coloring is by relative B-factor. Regions with the lowest B-factors are predicted to have the lowest mobility (*dark blue*); regions with the highest B-factors are predicted to have the highest mobility (*red*).

**Figure S7.**
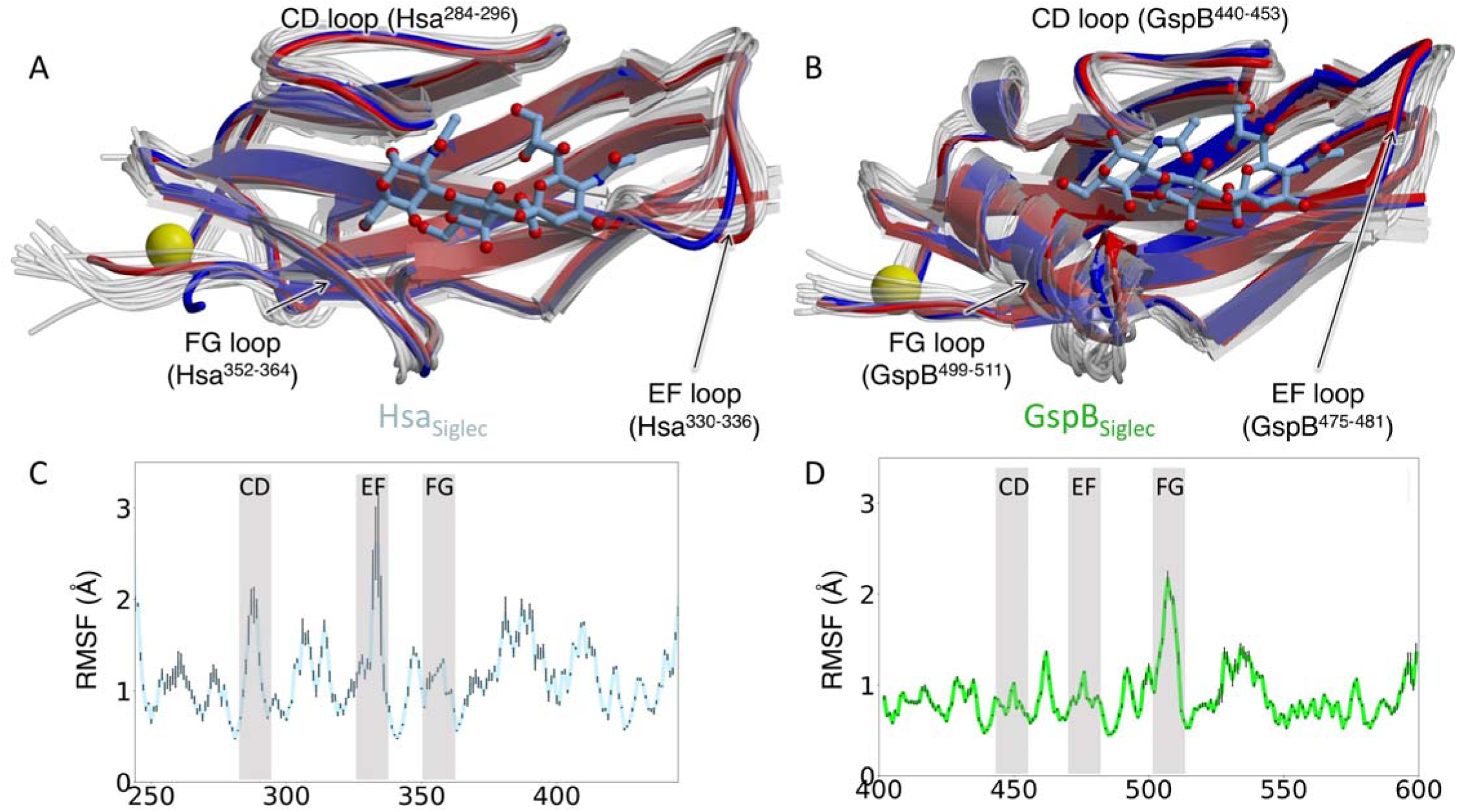
Conformational selection in SRR adhesins. A, B. Superposition of a representative subset of MD simulation snapshots (*translucent*) of A. Hsa and B. GspB onto the crystal structures determined in the presence (*blue*) and absence (*red*) of the sTa sialoglycan. MD simulations were performed on the adjacent Siglec and Unique domains; the Siglec domain was resected from the coordinates and is shown in isolation for clarity. MD simulations used structures determined in the absence of ligand as a starting point. C, D. Root mean square fluctuations (RMSF) of the Siglec domain of C. Hsa and D. GspB from the average position of the Cα atoms of each residue. Calculations were performed on the adjacent Siglec and Unique domains, with only the resected Siglec domain shown. Error bars correspond to the standard error over 3 independent simulations.

**Figure S8.**
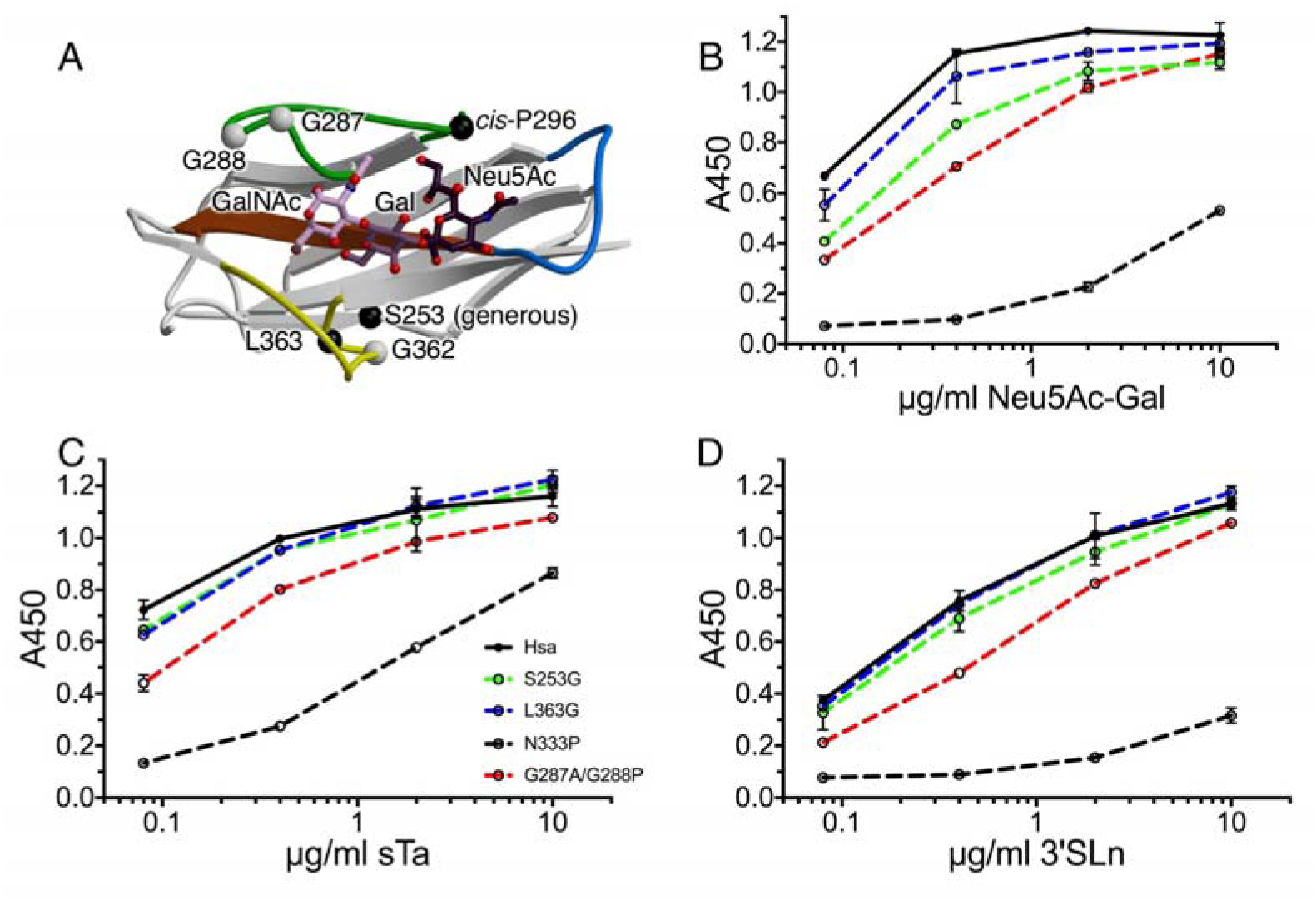
Impact of flexibility of sialoglycan binding by Hsa_BR_. A. Locations of variants within the Hsa_Siglec_ domain. As a note, Hsa^S253^ contains Ramachandran angles in the generously allowed region. B-D. Dose response curves of wild-type and variant GST-Hsa_Siglec+Unique_ (500 nM) immobilized in 96-well plates and binding to B. the Neu5Ac-Gal disaccharide, C. sialyl-T antigen, and D. 3’sialyl-*N*-acetyllactosamine. Biotinylated sialoglycan ligands were added at the indicated concentrations. Binding is reported as the mean ± standard deviation, with n = 2.

**Figure S9.**
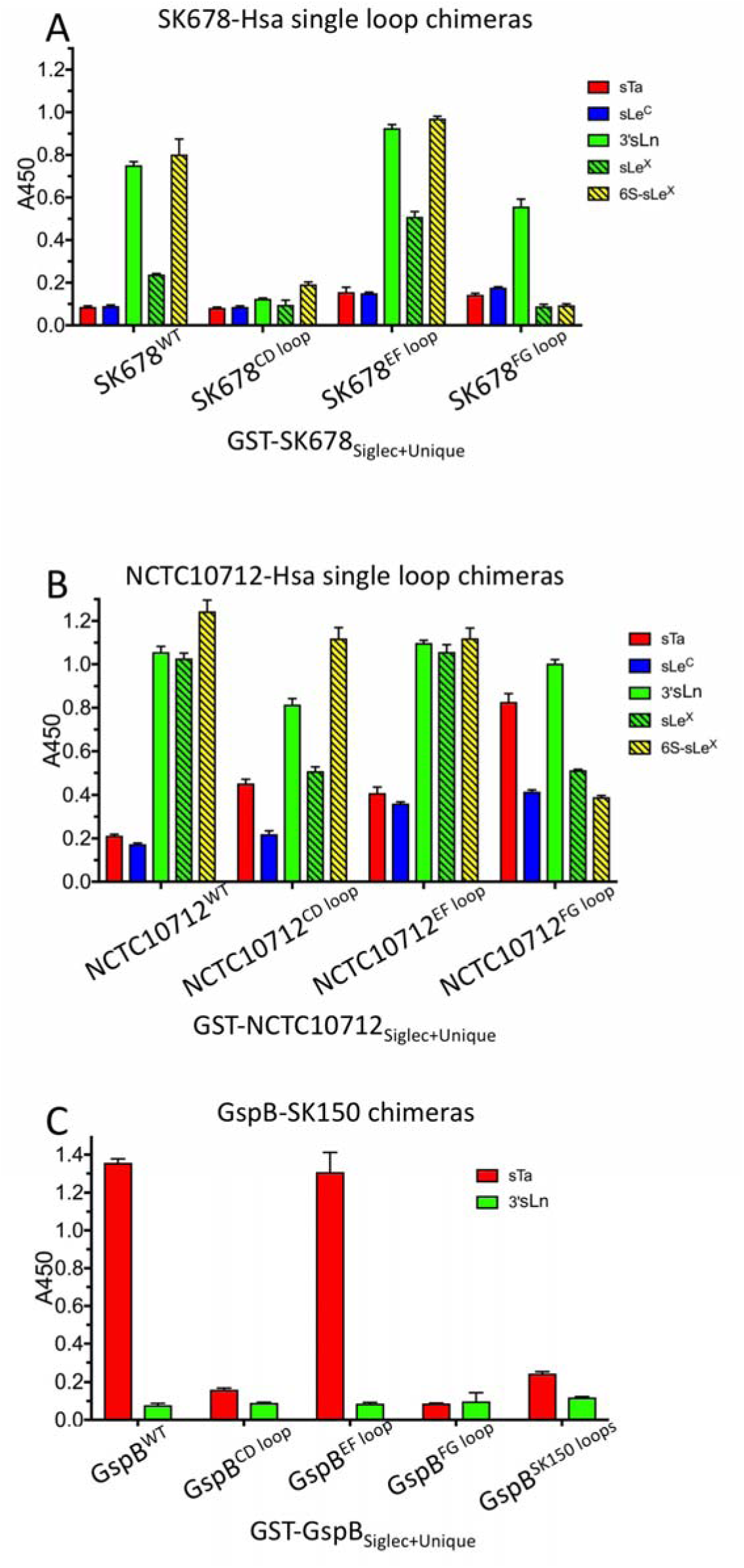
Chimeras of SRR adhesins. A. Binding of biotin-glycans (2 µg/ml) to GST-SK678_Siglec+Unique_ containing loops CD, EF, or FG of Hsa, substituted individually. Values correspond to the mean ± standard deviation, with n = 2 (wt) or n = 3 (variants). B. Binding of biotin-glycans (2 µg/ml) to GST-NCTC107128_Siglec+Unique_ containing loops CD, EF, or FG of Hsa, substituted individually. Values correspond to the mean ± standard deviation, with n = 2 (wt) or n = 3 (variants). C. Binding of biotin-glycans (1 µg/ml) to GST-GspB_Siglec+Unique_ containing loops CD, EF, or FG of SK150 substituted either individually or together. Values correspond to the mean ± standard deviation, with n = 3.

**Figure S10.**
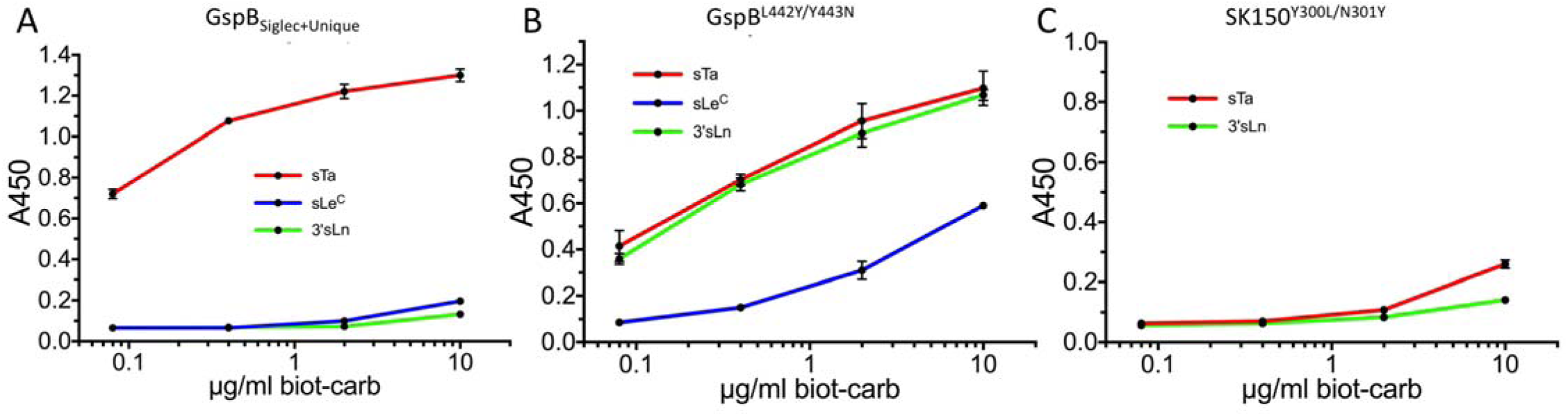
Mini-chimeragenesis of the GspB and SK150 adhesins. Dose-response curves of biotin-glycan binding to immobilized binding regions (500 nM). A. Wild-type GST-GspB_Siglec+Unique_ shows a binding preference for sTa. B. Mini-chimeragenesis with the SK150 adhesin was accomplished with the GST-GspB^L442Y/Y443N^ double mutant. The mini-chimera becomes more broadly selective by increasing the affinity for 3’sLn and sLe^C^. As a result, it exhibits binding selectivity more similar to wild-type GST-SK150_Siglec+Unique_ (see **Fig. S1**). C. The converse mini-chimeragenesis of SK150_Siglec+Unique_ exhibited reduced binding for sialoglycan ligands that bind most avidly to both wild-type GspB_Siglec+Unique_ and SK150_Siglec+Unique_.

**Figure S11.**
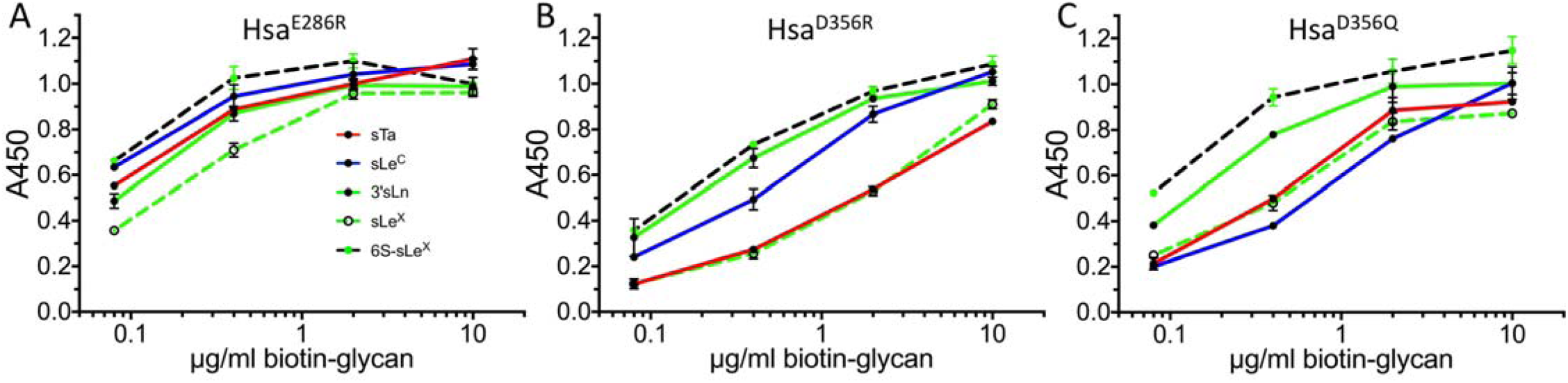
Binding characteristics of select GST-fused Hsa_Siglec+Unique_ variants. Dose-response curves of biotin-glycan binding to immobilized binding regions (500 nM). A. Hsa^E286R^, B. Hsa^D356R^, C. Hsa^D356Q^. Values are reported as the mean ± standard deviation, with n = 2.

